# Powerful and accurate case-control analysis of spatial molecular data

**DOI:** 10.1101/2025.02.07.637149

**Authors:** Yakir Reshef, Lakshay Sood, Michelle Curtis, Laurie Rumker, Daniel J. Stein, Mukta G. Palshikar, Saba Nayar, Andrew Filer, Anna Helena Jonsson, Ilya Korsunsky, Soumya Raychaudhuri

## Abstract

As spatial molecular data grow in scope and resolution, there is a pressing need to identify key spatial structures associated with disease. Current approaches typically make restrictive assumptions such as representing tissue regions by local abundances of manually typed, discrete cell types, or representing samples in terms of abundances of manually called, discrete spatial structures; this risks overlooking important signals. Here we introduce variational inference-based microniche analysis (VIMA), a method that combines deep learning with principled statistics to discover associated spatial features with greater flexibility and precision. VIMA trains an ensemble of variational autoencoders to extract numerical “fingerprints” from small tissue patches that capture their biological content. It uses these fingerprints to define a large number of data-dependent “microniches” – small, potentially overlapping groups of tissue patches with highly similar biology that span multiple samples. It then meta-analyzes across the autoencoders to identify microniches whose abundance correlates with case-control status while controlling for multiple testing. We show in simulations that VIMA is well calibrated. We then apply VIMA to spatial datasets spanning three different diseases and spatial modalities: a 7-marker immunofluorescence (IF) microscopy dataset in rheumatoid arthritis (RA), a 52-marker CO-Detection by indEXing (CODEX) dataset in ulcerative colitis (UC), and a 140-gene spatial transcriptomics dataset in dementia. In each case, we recapitulate known biology and identify novel spatial features of disease that were not discoverable with current state-of-the-art methods.

## Main

Spatial profiling of tissue samples has been a foundational tool in biology starting from the discovery of capillaries with the microscope in the 1600s^1^. With the introduction of haematoxylin and eosin (H&E) staining in the late 1800s^2^, spatial *molecular* techniques began to provide additional insight and are now a pillar of both clinical medicine and biomedical research^3–6^. In recent years, the scale and richness of spatial molecular modalities have dramatically increased. Highly multiplexed protein modalities such as imaging mass cytometry^7^ and CODEX^8^ have enabled the spatial measurement of scores of proteins, and transcript-based modalities such as MERFISH^9^, Xenium^10^, and others^11–14^ can now profile thousands of genes at high spatial resolution.

However, the very richness of modern spatial data has made systematic research challenging. Historically, spatial data have been analyzed through manual inspection and the development of highly detailed, bespoke assessment systems. For example, in rheumatoid arthritis, synovial inflammation is graded using a complex scoring system that relies on qualitative assessments made by a pathologist^15^, and tumors are characterized using assessments of cancer cell morphology and of lymphocyte arrangements and proportions in specific tumor microenvironments^16,17^. These assessment systems limit discoveries to those that fit the paradigms under which they were established and scale poorly as we assay more genes and proteins.

Thus, the development of new computational methods for high-resolution annotation of tissue structures in spatial molecular data has become an active area of research. Modern methods based on local abundances of different cell types^17–19^, spatially constrained clustering^20^ and aggregation^21,22^, spatial factorization^23,24^, graph neural networks^25–27^, and others^28–30^ have enabled identification of granular tissue structures. More recent work has begun to explore the intuition that neural network architectures from image analysis can be built upon to provide powerful and flexible tissue representations^31–33^.

Here, we focus specifically on the question of statistical case-control analysis in spatial data: given a spatial dataset with multiple case and control samples, are there patterns that are statistically significantly different between cases and controls, after multiple testing correction? One simple strategy for addressing this question is to annotate the samples in such a dataset with a modern tissue annotation method, use the annotations to cluster regions of tissue with similar biology, and then to test for differential abundance of the clusters in cases versus controls. However, while this approach has been used successfully to find case-control associations^17,19,21,22,32^, it may be under-powered if the tissue annotation is not sufficiently expressive, if the case-control differences are obfuscated by the hard clustering step, or if the tissue annotation is too sensitive to sample-specific idiosyncrasies.

To address this challenge, we introduce Variational Inference-based Microniche Analysis (VIMA), a method for statistical case-control analysis of molecular spatial data. VIMA uses neural network-based techniques from image processing to learn compact, informative “fingerprints” for each of many small tissue patches in a spatial dataset, similarly to some recent methods^32,33^. However, VIMA introduces three innovations. First, it uses a conditional variational autoencoder (cVAE) architecture designed to learn fingerprints that are minimally influenced by sample- or batch-specific effects, thereby enhancing sensitivity to true case-control signals. Second, VIMA leverages the stochasticity of neural network optimization by training an ensemble of cVAEs rather than just one, resulting in multiple representations that can capture complementary aspects of the same dataset while also increasing replicability. Third, rather than using the fingerprints to cluster the tissue patches into discrete groups, VIMA instead defines many small, potentially overlapping groups of highly similar tissue patches, which we term *microniches*, and quantifies the abundance of each microniche in each sample. VIMA then performs statistical tests to identify non-trivial correlations between the abundances of the microniches and case-control status, meta-analyzing across the ensemble of cVAEs.

VIMA produces both a global statistical test for the presence of any case-control difference in the spatial data as well as a local statistical test identifying the set of specific patches explaining that difference at a given false discovery rate threshold. VIMA can be applied to any spatially resolved molecular technology, is well powered even at the modest sample sizes typical of research cohorts, and avoids traditional, parameter-intensive preprocessing steps such as cell segmentation, clustering of cells into discrete cell types, or clustering of tissue regions into discrete groups. VIMA produces properly calibrated P-values and so can be used for statistical hypothesis testing, a fact that we confirm in null simulations.

We apply VIMA to three spatial molecular case-control datasets containing four case-control signals and generated with three different technologies in rheumatoid arthritis (7-marker immunofluorescence microscopy), ulcerative colitis (52-marker CODEX), and dementia (140-gene MERFISH). In each case, VIMA identifies spatial features that differ significantly between cases and controls and that both recapitulate known biology and provide new insights. We also benchmark VIMA against seven state-of-the-art methods and find that most of VIMA’s signals are not detectable with these methods. Finally, we show using an ablation study that each novel component of VIMA is necessary for its performance.

## Results

### Overview of methods

VIMA is based on three central concepts. First, we use a deep learning architecture common in image analysis – augmented to account for sample-specific spatial artifacts – to represent each tissue patch in a dataset with a numerical “fingerprint” summarizing its essential biology. Second, we take advantage of the stochasticity of neural network training to generate ten sets of these fingerprints, increasing our sensitivity to underlying biology while simultaneously improving replicability. Third, rather than using the fingerprints to cluster tissue patches into discrete, non-overlapping groups, we instead define many small, overlapping groups of highly similar tissue patches called “microniches” that form the basic unit of association testing.

VIMA takes as input 1) a multi-sample spatial molecular dataset that profiles protein levels, transcript levels, stain intensities, or some combination of these, and 2) a table of sample-level metadata containing case-control status and any other relevant covariates such as patient age or sex. VIMA’s output consists of 1) a tensor of samples by microniches by autoencoders called the microniche abundance tensor (MAT) that summarizes the abundance patterns of different microniches across the samples, 2) a global P-value indicating the statistical significance of the aggregate spatial differences between case samples and control samples, and 3) a subset of tissue patches whose microniches drive this association at a specified false discovery rate threshold, with a directional effect size for each patch. VIMA’s results can be represented via a consensus nearest neighbor graph of tissue patches that can be visualized with UMAP, with similar tissue patches showing up near each other in UMAP space.

To create the patch fingerprints, VIMA rasterizes each sample into pixels, performs principal components analysis (PCA) to reduce the number of markers per pixel to a smaller set of “meta-markers”, and integrates these meta-markers using Harmony^34^ to remove non-spatial batch effects and sample-specific artifacts (**Methods**). VIMA then transforms each sample in the dataset into a collection of partially overlapping, square tissue patches (**Fig. 1A**) that are 400×400um by default. It then uses the tissue patches to train an ensemble of ten conditional variational autoencoders^35^ (cVAEs) with identical ResNet18-style architectures^36^ but different initial weights; each cVAE extracts from every tissue patch a numerical latent representation (the “fingerprint”; **Fig. 1B**) while explicitly being incentivized to remove the effects of sample ID on the fingerprint (**Methods**).

**Figure 1:**
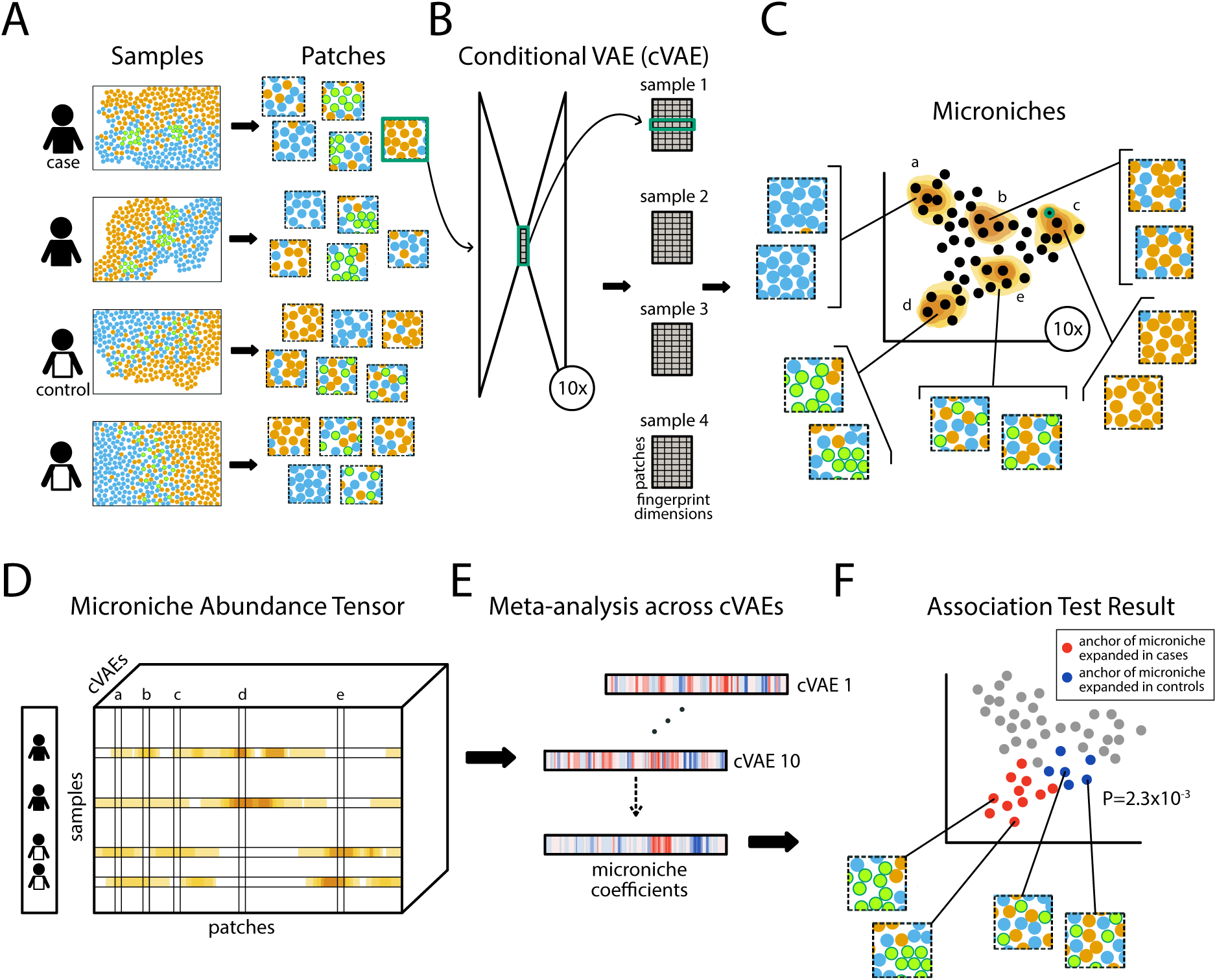
A schematic depiction of VIMA. **A)** VIMA generates a large number of small, square, partially overlapping tissue patches from each sample. **B)** VIMA trains ten identical conditional variational autoencoders (cVAEs) with different initial conditions on all the tissue patches to learn ten low-dimensional vector representations (“fingerprints”) of each patch that summarize its biological contents, such that similar patches have similar fingerprints. **C)** VIMA uses each set of fingerprints to build a nearest-neighbor graph of patches and then defines, for each graph, a microniche anchored at each patch to which every other patch in the graph belongs in proportion to how close it is to the anchor patch in the graph. **D)** VIMA counts the fraction of patches in each sample belonging to each microniche and represents this information as a samples-by-microniches-by-cVAEs tensor called the microniche abundance tensor, alongside the sample-level phenotype. **E)** VIMA correlates each micronice in the MAT with the phenotype, meta-analyses across the 10 cVAEs to assess which patches have significant correlations after accounting for multiple hypothesis testing and correlation among tests. We visualize the result by coloring each patch in a consensus UMAP according to the meta-analyzed correlation of the abundance of its respective microniches to the phenotype, known as its “microniche coefficient”. Patches with non-significant microniche coefficients are colored in gray. The P-value overlaid on the plot is a global P-value that reflects the overall significance of the case-control association.

To define microniches, VIMA builds on our previous work in single-cell analysis^37^. For each of the ten cVAEs, it first constructs a nearest-neighbor graph of the patch fingerprints produced by that cVAE (**Fig. 1C**). It then defines a single microniche anchored at each patch *p*: every other patch *p’* belongs to that microniche according to the probability that a random walk in the graph from *p’* will arrive at *p* (**Methods** and **Fig. 1C**). VIMA uses the microniches to compute a microniche abundance tensor (MAT) whose *n,p,k*-th entry is the fraction of patches from sample *n* in the microniche anchored at patch *p* that was defined using the *k*-th cVAE (**Fig. 1D**). VIMA meta-analyzes the MAT across the ten cVAEs (**Fig. 1E**) to perform both a “global” and a “local” association test. The global test tests for the simple presence of any association between the *pk* microniches in the MAT and case-control status; the local test tests each patch to determine whether any of the ten microniches anchored at that patch are driving the association (**Fig. 1F**).

To perform the local test, VIMA computes the correlation between the abundance of each microniche and case-control status and then uses an adaptive weighted averaging scheme to meta-analyze the ten correlations corresponding to each patch into a single correlation called the “microniche coefficient” of that patch (**Fig. 1E**). It then estimates empirical false discovery rates (FDRs) for each microniche coefficient by permuting sample labels and re-computing the same statistics to obtain null distributions (**Fig. 1F**). To perform the global test, the per-microniche correlations are combined in a data-dependent way into a single aggregate test statistic across the entire MAT whose significance is again estimated using null permutations (**Fig. 1F**). In both the global and local tests, VIMA can control for sample-level confounders, such as demographic variables and technical parameters (**Methods**). We confirmed VIMA’s type I error control in simulations (**Supplementary Fig. 1**).

VIMA requires no pre-training, making it simple to apply out-of-the-box to new datasets and new technologies. It requires minimal parameter tuning and has favorable runtime properties: for the three datasets analyzed in this paper containing S=27, S=42, and S=75 samples, respectively, the total runtime of the entire VIMA workflow including preprocessing was just ∼2 hours, ∼2 hours, and ∼8 hours, respectively, on a single GPU. We have released open-source software implementing the method.

### Identifying spatial features of rheumatoid arthritis subtypes using immunofluorescence microscopy data

Rheumatoid arthritis is an inflammatory arthritis affecting up to 1% of the population in which immune infiltration alters the synovium of the joints, leading to tissue destruction^38^. We applied VIMA to a 7-marker IF microscopy dataset of S=27 synovial biopsies from N=22 rheumatoid arthritis patients collected by the Accelerating Medicines Partnership Rheumatoid Arthritis/Systemic Lupus Erythematosus (AMP RA/SLE) consortium^39^. (See **Supplementary Table 1** for the list of markers profiled.) The original analysis of this dataset used a superset of this cohort (N=70) in which ∼20,000 genes were profiled with single-cell RNA-seq to assign each sample to one of six different “cell-type abundance phenotypes” (CTAPs). Due to the limited size of the smaller cohort that was profiled with IF microscopy, we grouped the CTAPs here into three “stromal” types, defined by high abundance of fibroblasts, and three “immune” types, defined by a high abundance of T cells, B cells, and myeloid cells. This grouping was motivated by the emerging role in RA of fibroblasts^40,41^, which have recently been recognized as correlating with poor response to current treatment modalities^39^.

Before applying VIMA’s local and global tests for association to stromal vs immune status, we first examined the properties of VIMA’s patch fingerprints and their associated microniches. The VIMA autoencoders reconstructed the tissue patches with an average mean-squared error (MSE) of 0.57. This is a substantial improvement over the MSE of 1.00 that results from guessing the dataset-wide average for all pixels and the MSE of 0.75 that results from guessing the average expression profile of each patch. **Fig. 2A** shows an example of a tissue patch from the dataset together with its reconstruction based on a single patch fingerprint from one cVAE. We found that, in addition to enabling accurate reconstruction, the patch fingerprints effectively accounted for sample-specific idiosyncracies. We quantified local sample mixing by calculating the perplexity across samples of each patch’s fingerprint-based nearest neighbors. The VIMA-based fingerprints produced significantly higher perplexities than those generated by other methods, including other neural-network based methods (**Fig. 2B**; **Methods**). (Due to the lack of cell segmentation in this dataset, we were only able here to benchmark three of the seven methods in our benchmarking suite, as the others require segmented cells for their input. We profile additional methods in the subsequent two datasets.)

**Figure 2:**
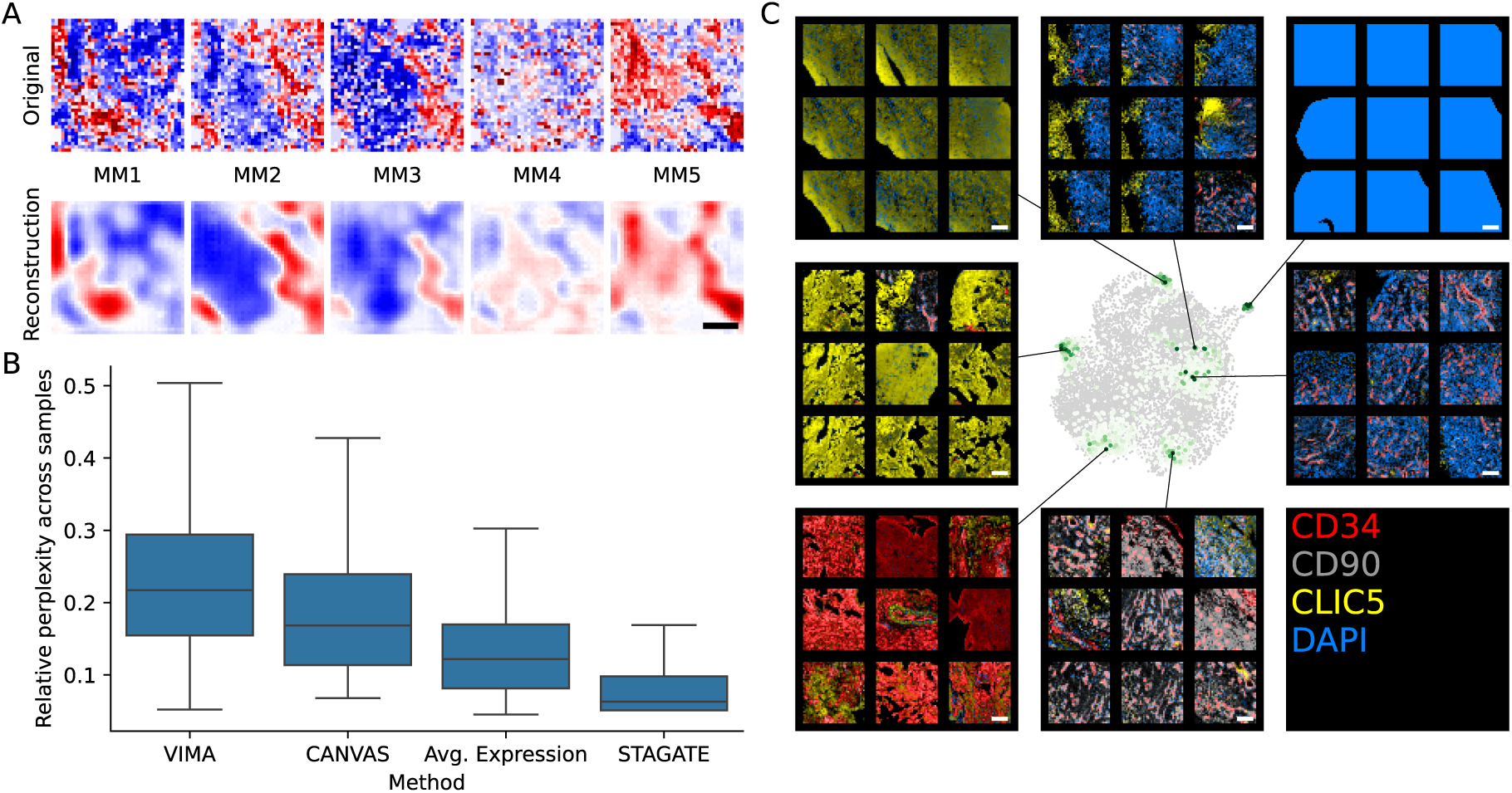
Basic properties of VIMA’s microniches in the RA dataset. A) Direct comparison of an example tissue patch (top row) against the reconstruction provided by one of VIMA’s autoencoders (bottom row). For each of the five meta-markers (MMs), we depict the marker value across the pixels in the original and the reconstructed patch. The meta-marker used to color pixels in each pair of pictures is indicated by the labels MM1, …, MM5.B) Comparison of sample-level integration across methods. For each method, we computed a relative perplexity score reflecting how evenly each spatial unit considered by that method connects in the nearest neighbor graph to spatial units from a diversity of samples (Methods). We show the distributions of these scores for all spatial units considered by each method. C) A UMAP of all the patches in the dataset generated using one of VIMA’s sets of patch fingerprints, together with randomly selected patches drawn from 7 example microniches in the dataset; these microniches are characterized by (clockwise from bottom-left): high CD34 intensity with interspersed CLIC5 signal, high CLIC5 intensity with minimal CD34 signal, high CLIC5 intensity but with a blurrier texture and interspersed DAPI signal, a vertical boundary between CLIC5-high/DAPI-low pixels and CLIC5-low/DAPI-high pixels, a DAPI staining artifact, DAPI-high infiltrate surrounding CD34-high regions that are likely blood vessels, and perivascular CD90-high pixels. All scale bars represent 100um.

On visual inspection, the microniches generated by VIMA’s autoencoders appear to exhibit consistent biological features (**Fig. 2C**). For example, examining the output of one arbitrarily selected autoencoder, we observed microniches containing: broad CD34-high staining with small interspersed CLIC5-high areas, likely representing CD34+/CD90-sublining fibroblasts surrounding small collections of lining fibroblasts; large areas of CLIC5-high staining with minimal CD34 signal, indicative of regions of lining fibroblasts; CLIC5-high areas but with a blurrier staining texture and interspersed DAPI signal; a vertical boundary between CLIC5-high regions of lining fibroblasts and CLIC5-low/DAPI-high regions; DAPI saturation that likely represents staining artifact; DAPI-high infiltrate surrounding CD34-high regions that are likely blood vessels; and perivascular CD90-high staining that represents either pericytes or CD90+ lining fibroblasts surrounding blood vessels. These microniches reinforce the potential of VIMA’s cVAE’s to aggregate together areas with concordant tissue features across multiple samples in a dataset.

We next asked whether VIMA could “re-discover” the stromal vs immune distinction found in the larger single-cell cohort in this smaller dataset, without being given this information upfront. To do this, we applied PCA to the MAT and examined the sample loadings along the first two PCs. Indeed, these two PCs nearly perfectly separate the stromal samples from the immune samples (**Fig. 3A**). Thus, although this dataset has only 7 markers from 22 donors, rather than 20,000 genes from 70 donors used in discovery, VIMA is able to effectively leverage the power of spatial information to clearly reveal this important source of biological variation across the samples.

**Figure 3:**
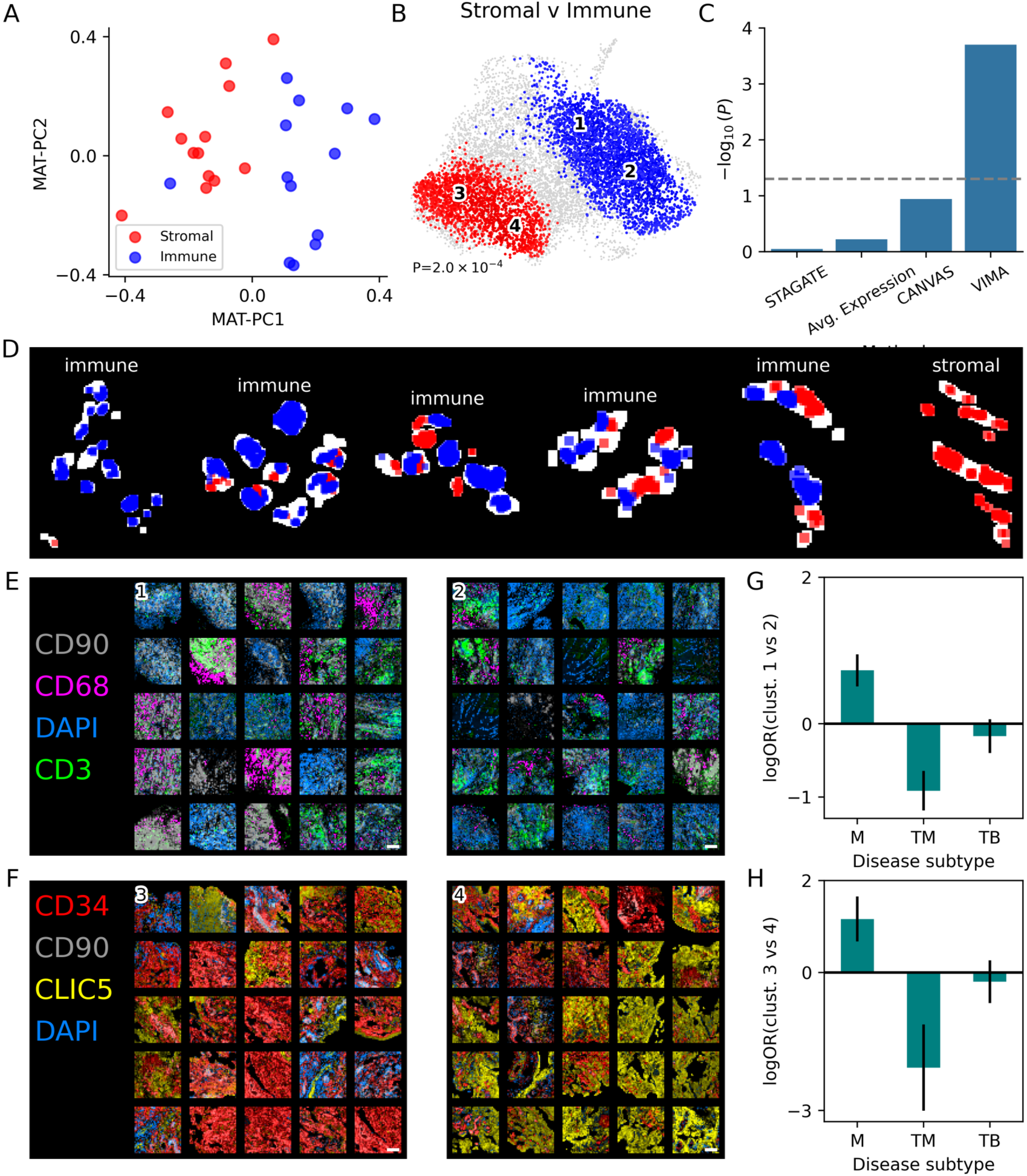
VIMA analysis of disease subtypes in the RA dataset. A) plot of MAT-PC1vs MAT-PC2, with each dot representing a sample and samples colored by RA subtype as determined using a larger scRNA-seq dataset (Methods). B) Results of VIMA case-control analysis for stromal vs immune subtypes: a UMAP of all the patches in the dataset is generated using the patch fingerprints, and the patches whose microniche coefficients pass the 10% FDR threshold are colored in proportion to the meta-analyzed correlation between the abundances of their microniches and stromal status, with red denoting positive correlations and blue denoting negative correlations. The centroids of the two subclusters of each of the positive and negatively associated patches, respectively, are indicated with numbers. The global P-value for the association test is overlaid. C) Comparison of VIMA’s global P-value for the stromal vs immune association to those obtained with tissue representations produced by other methods (Methods). D) A subset of 6samples from the dataset exemplifying both spatial homogeneity in local stromal vs immune status within a sample (the leftmost and rightmost samples) and spatial heterogeneity in local stromal vs immune status within a sample (middle samples), with disease subtype indicated above each sample. Each patch in each sample is colored red if it is stromal-associated by VIMA or blue if it is immune-associated. E) Randomly selected patches from each of the two clusters of immune-associated patches. F) Randomly selected patches from each of the two clusters of stromal-associated patches. G) Log-odds ratios among the immune-associated patches for belonging to cluster 1 (positive log-odds) vs cluster 2 (negative log-odds) as a function of the finer-grained disease subtypes in the dataset, with 95% confidence intervals; M: myeloid-predominant, TM: T cell/myeloid-predominant, TB: T cell/B cell-predominant; results for other subtypes are shown in Supplementary Fig. 5. H) Log-odds ratios among the stromal-associated patches for belonging to cluster 3 (positive log-odds) vs cluster 4 (negative log-odds) as a function of the same finer-grained disease subtypes. Scale bars represent 100um throughout.

We then applied VIMA’s local and global tests to formally test for differences between the stromal and immune samples. VIMA identified a strong case-control difference (P=2.0e-4) with 5,222 of 10,345 patches significant at FDR 10% (**Fig. 3B**), consistent with the separation of stromal from immune samples in the unsupervised analyses above. Reassuringly, the stromal-associated patches were higher in CLIC5 (a marker of lining fibroblasts) and the immune-associated patches were higher in CD3 (T cells), CD68 (macrophages), and DAPI (nucleated cells; **Supplementary Fig. 2**). VIMA therefore is able to recapitulate the biology we expect, validating the biological interpretability of its approach. In contrast, we were unable to detect a statistically significant difference between the stromal and immune samples using discrete tissue annotations generated by the other spatial methods that we benchmarked on this dataset (**Fig. 3C**; **Methods**).

The spatial nature of our results enabled us to examine for the first time the spatial distribution of areas of stromal and immune biology *within* each sample. To our surprise, we found that even though each sample received just one overall label from the non-spatial single-cell data, samples with the immune label in fact often contained both stromal-like regions and immune-like regions that are spatially segregated (**Fig. 3D**). The cells of the synovium therefore form spatially organized communities in specific regions of tissue that have distinct types of biology and are not adequately captured by sample-wide cell-type abundances. This spatial characterization advances our understanding of how key cell types in the synovium work to create disease states. It also challenges the concept of a single disease subtype per patient by demonstrating the biological heterogeneity that can exist within even a single patient, with implications for precision medicine in this disease.

We also found clear biological heterogeneity among the tissue patches associated with stromal and immune status, respectively. For example, the immune-associated patches clustered into two subtypes (**Fig. 3E**). The first is highly cellular, characterized by strong DAPI staining intermixed with a T-cell infiltrate (**Fig. 3E**, cluster 1; **Supplementary Fig. 3**). The second is characterized by spatial proximity of areas high in the macrophage marker CD68 and areas high in the sublining fibroblast marker CD90, with some T cell aggregates (**Fig. 3E**, cluster 2; **Supplementary Fig. 3**). Similarly, the stromal-associated patches also clustered into two subtypes (**Fig. 3F**). The first is characterized by strong staining of CD34, a marker of sublining fibroblasts and endothelial cells (**Fig. 3F**, cluster 3; **Supplementary Fig. 4**). The second is characterized by strong staining of CLIC5, a marker of lining fibroblasts (**Fig. 3F**, cluster 4; **Supplementary Fig. 4**).

We found strong associations between these clusters and 2 of the 6 more granular, original disease subtypes (the CTAPs) in this cohort. Of the two immune-associated clusters, cluster 1 was more prevalent than cluster 2 in the “myeloid” (M) CTAP (log-odds-ratio 0.73, 95% confidence interval [0.51,0.95]) and less prevalent than cluster 2 in the “T-cell/myeloid” (TM) CTAP (log-odds-ratio −0.91, 95% confidence interval [−1.18,-0.65]; **Fig. 3G**). And of the two stromal-associated clusters, cluster 3 was more prevalent than cluster 4 in samples from the M CTAP (log-odds-ratio 1.16, 95% confidence interval [0.67,1.65]) and less prevalent than cluster 4 in samples from the TM CTAP (log-odds-ratio −2.07, 95% confidence interval [−3.01,-1.13]; **Fig. 3H**). (Other CTAPs had no significant enrichments; **Supplementary Fig. 5**.) Overall, these findings suggest that the M CTAP may be characterized by vascularization, macrophage/sublining-fibroblast interactions, and separate T-cell aggregates, whereas the TM CTAP may be characterized by an expanded lining and an as-yet uncharacterized immune infiltrate with a relationship to T cell activation. Therefore, although these two CTAPs have quite similar sample-wide cell-type abundance profiles in the single-cell data (myeloid-predominant vs T cell/myeloid predominant), each appears to have quite distinct underlying tissue biology. VIMA’s ability to spatially characterize the tissue patches in this dataset suggests that spatial profiling with a larger marker set may unlock *** a clearer elucidation of the biology driving these disease subtypes.

### Identifying effects of TNF inhibition on lymphoid aggregates in ulcerative colitis using CODEX data

Ulcerative colitis is a chronic inflammatory disease characterized by inflammation and ulceration of the colonic mucosa that leads to symptoms such as diarrhea, abdominal pain, and rectal bleeding^42^. In the last decade, inhibition of tumor necrosis factor (TNF) has emerged as a cornerstone of treatment^42^. However, not all patients respond to TNF inhibition, and the molecular determinants of this heterogeneity are not known^43^. To identify the spatial consequences of TNF inhibition in the colon, we applied VIMA to a 52-marker CODEX^8^ dataset of S=42 colonic biopsies from N=34 patients, 29 of whom had ulcerative colitis^19^ and the rest of whom were healthy. (See **Supplementary Table 2** for the list of markers profiled.)

We first used VIMA for the relatively easier task of distinguishing UC samples from healthy samples and found a strong association (global P= 1.7e-3) with 11,839 of 21,359 significant patches at FDR 10%, of which 4,146 were UC-associated and 7,693 were healthy-associated (**Fig. 4A**). We found that the UC-associated patches tended to lie deep to the epithelial layers, consistent with known immune-cell infiltration of these layers in ulcerative colitis^44,45^ (**Fig. 4B**). As expected with an inflammatory disease, VIMA found that the UC-associated patches had significantly higher average levels of immune cell markers such as CD3 (T cells), CD19 (B cells), and CD45 (most immune cells; **Fig. 4C**). Unlike with the RA case-control analysis, the UC-versus-healthy signal was detectable using discrete tissue annotations generated by two other spatial methods that we benchmarked: UTAG^21^ and a baseline of patchwide average expression profile after pixel-level Harmony^34^ (**Fig. 4D**). However, VIMA detected the signal with a substantially higher level of significance. In addition, VIMA was the only method that detected UC-associated patches and not just healthy-associated patches, a crucial prerequisite for characterizing the effect of TNF inhibition on UC-affected portions of the colon (**Fig. 4E**). In the remainder of our analysis, we restricted our attention to the 4,146 patches that VIMA detected as UC-associated.

**Figure 4:**
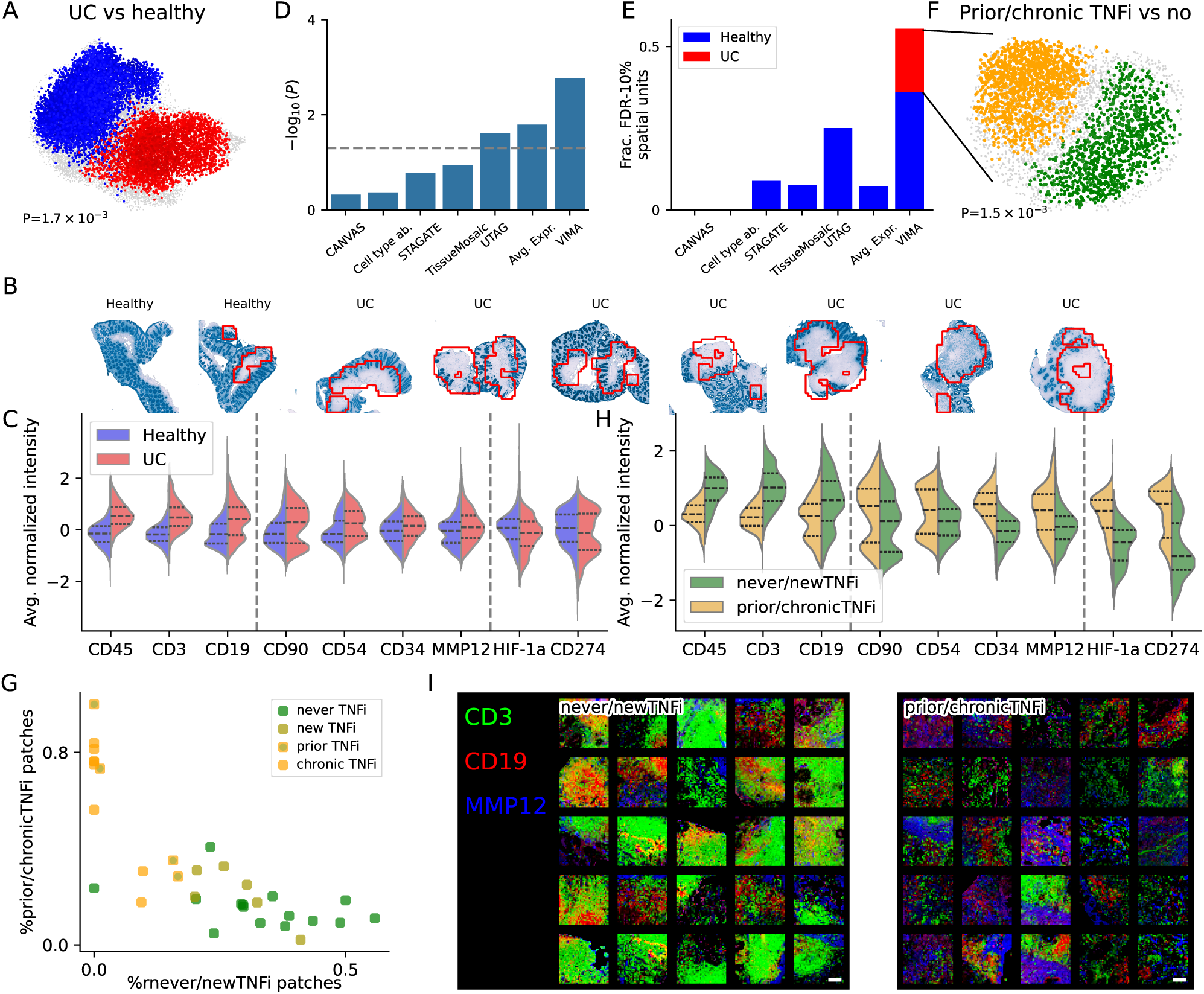
VIMA analysis of UC dataset. **A)** Results of VIMA case-control analysis for UC versus healthy: a UMAP of all the patches in the dataset is generated using the patch fingerprints, and the patches whose microniche coefficients pass the 10% FDR threshold are colored in proportion to the meta-analyzed correlation between the abundances of their microniches and UC status, with red denoting positive correlations and blue denoting negative correlations. The global P-value for the association test is overlaid. **B)** Selected samples shown with cytokeratin staining in blue and contiguous areas of UC-associated patches outlined in red. **C)** Distributions of average intensity per patch of selected markers among UC-associated and healthy-associated patches. The selected markers are grouped using dotted lines according to the relationship of their directions of effect to those in panel (H). **D)** Comparison of VIMA’s global P-value for the UC vs healthy association to those obtained with tissue representations produced by other methods (**Methods**). **E)** Comparison across methods of the number of spatial units that are significantly associated with UC (red) and healthy (blue) status at FDR 10%. **F)** VIMA analysis within UC-associated patches comparing the samples with prior or chronic TNF inhibition (prior/chronicTNFi) to those with no or recent TNF inhibition (never/newTNFi), with gold denoting positive correlation to prior/chronicTNFi status and green denoting negative correlation. The global P-value for the association test is overlaid. **G)** Comparison of fraction of prior/chronicTNFi-associated vs never/newTNFi-associated patches in each sample. For each sample, the percent of patches in that sample that are significantly associated with prior/chronicTNFi is plotted against the percent of patches in that sample significantly associated with never/newTNFi. Each sample is then colored by its detailed TNFi status. **H)** Distributions of average intensity per patch of selected markers among prior/chronicTNFi-associated and never/newTNFi-associated patches. The selected markers are grouped using dotted lines as follows: markers that are higher in UC compared to control and in never/newTNFi compared to prior/chronicTNFi (left), markers that are higher in UC compared to control but lower in never/newTNFi compared to prior/chronicTNFi (middle), and markers that are lower in UC compared to control and lower in never/newTNFi compared to prior/chronicTNFi (right). **I)** Randomly selected patches from the never/newTNFi-associated patches (left) and the prior/chronicTNFi-associated patches (right).

To study the effects of TNF inhibition on UC biology, we used VIMA to conduct a case-control analysis only among UC-associated patches comparing UC patients treated with TNF inhibition in the past (“prior/chronicTNFi”) to UC patients with either no TNF inhibition at all or recent TNF inhibition only (“never/newTNFi”; **Methods**). Strikingly, we found a strong association (global P=1.5e-3) with 2,307 of the 4,146 patches significant at FDR 10% (**Fig. 4F**). Moreover, the proportion of patches in each sample that were significantly prior/chronicTNFi-associated versus never/noTNFi-associated strongly stratified samples by their more granular TNF status in a dose response-like fashion: we observed a gradient with patients treated with longstanding TNF inhibition on one end, followed by patients with prior but not current TNF inhibition, then patients with recent initiation of TNF inhibition, and finally patients who have never been on TNF inhibition (**Fig. 4G**).

The prior/chronicTNFi-associated patches had some features in common with healthy tissue and other features in common with UC tissue. For example, they had lower levels of traditional B- and T-cell markers, similar to the healthy-associated patches, but they also had higher levels of fibroblast, macrophage, and mesenchymal markers such as CD54, CD90, CD34, and MMP12 (**Fig. 4H**), similar to the UC-associated patches (**Fig. 4C**). We verified these differences at the level of microniche coefficients as well (**Supplementary Fig. 6**). We next examined the spatial relationships among the markers *within* the associated tissue patches. We found that many of the never/newTNFi-associated patches were characterized by B-cell aggregates with surrounding T-cells, consistent with lymphoid aggregates, while the chronic/priorTNFi-associated patches showed a spatially disorganized pattern of mesenchymal markers intermixed with scant B- and T-cells (**Fig. 4I**). Overall, these results suggest that TNF inhibition strongly affects colonic tissue architecture, perhaps even persistently after treatment, and that TNF inhibition specifically reduces lymphoid aggregates while upregulating mesenchymal and fibroblast activity.

Our results build on and strengthen observations made in the index analysis of this dataset^19^. In that analysis, the authors used a detailed, cell-type based local averaging workflow requiring segmentation of each sample into cells, clustering of the cells into cell types, and clustering of spatial regions on the basis of cell-type abundance into “cellular neighborhoods”. In one specific subgroup of UC patients – those with intermediate-grade colonic inflammation – this analysis strategy revealed a modest negative association (P<0.01) between treatment with TNF inhibition and the presence of one of the 10 cellular neighborhoods annotated as containing lymphoid aggregates. While suggestive, this association was found in one of several possible patient subsets and was not corrected for multiple hypothesis testing. In contrast, VIMA directly identifies a statistically robust reduction in lymphoid structures in TNF-treated patients, without the need for segmentation, cell-type annotation, and patient subset selection, providing more definitive evidence that TNF inhibition disrupts lymphoid aggregation in colonic tissue.

### Identifying dementia-associated tissue structures using spatial transcriptomics data

Alzheimer’s disease is the most common cause of dementia in older patients^46^. Efforts to characterize the molecular signatures of Alzheimer’s dementia using single-cell RNA-seq have identified key cell populations whose aggregate abundance is either increased or decreased in Alzheimer’s patients^47^. However, the question of which multi-cellular spatial structures are characteristic of Alzheimer’s dementia, and of dementia more broadly, has been difficult to study with non-spatial single-cell technologies. We applied VIMA to a 140-gene MERFISH dataset consisting of S=75 post-mortem medial temporal gyrus samples from N=27 patients, 15 of whom had either Alzheimer’s dementia or an Alzheimers-related dementia^47^, with a total of 516 million transcripts profiled. (See **Supplementary Table 3** for the list of genes profiled.)

In our analysis, VIMA detected a significant association to dementia status with many significant patches, despite the lower sample size of the spatial data (global P=5.4e-3; 27,072 out of 70,898 patches significant at FDR 5%; **Fig. 5A**). Moreover, the association appears quite strong: the percent of dementia-associated vs control-associated patches in each sample nearly separates dementia samples from control samples (**Fig. 5B**). In contrast, we were unable to detect this association using annotations generated by the other spatial methods that we benchmarked (**Fig. 5C**). To quantify the power gain of a spatial analysis with VIMA as opposed to a case-control analysis based only on non-spatial single-cell data at the same sample size, we also conducted a case-control analysis just using the sample-wide cell-type proportions for each sample; this analysis was also null, suggesting that a non-spatial, cell type-based single-cell analysis would need a larger sample size to detect this association (**Supplementary Fig. 7**).

**Figure 5:**
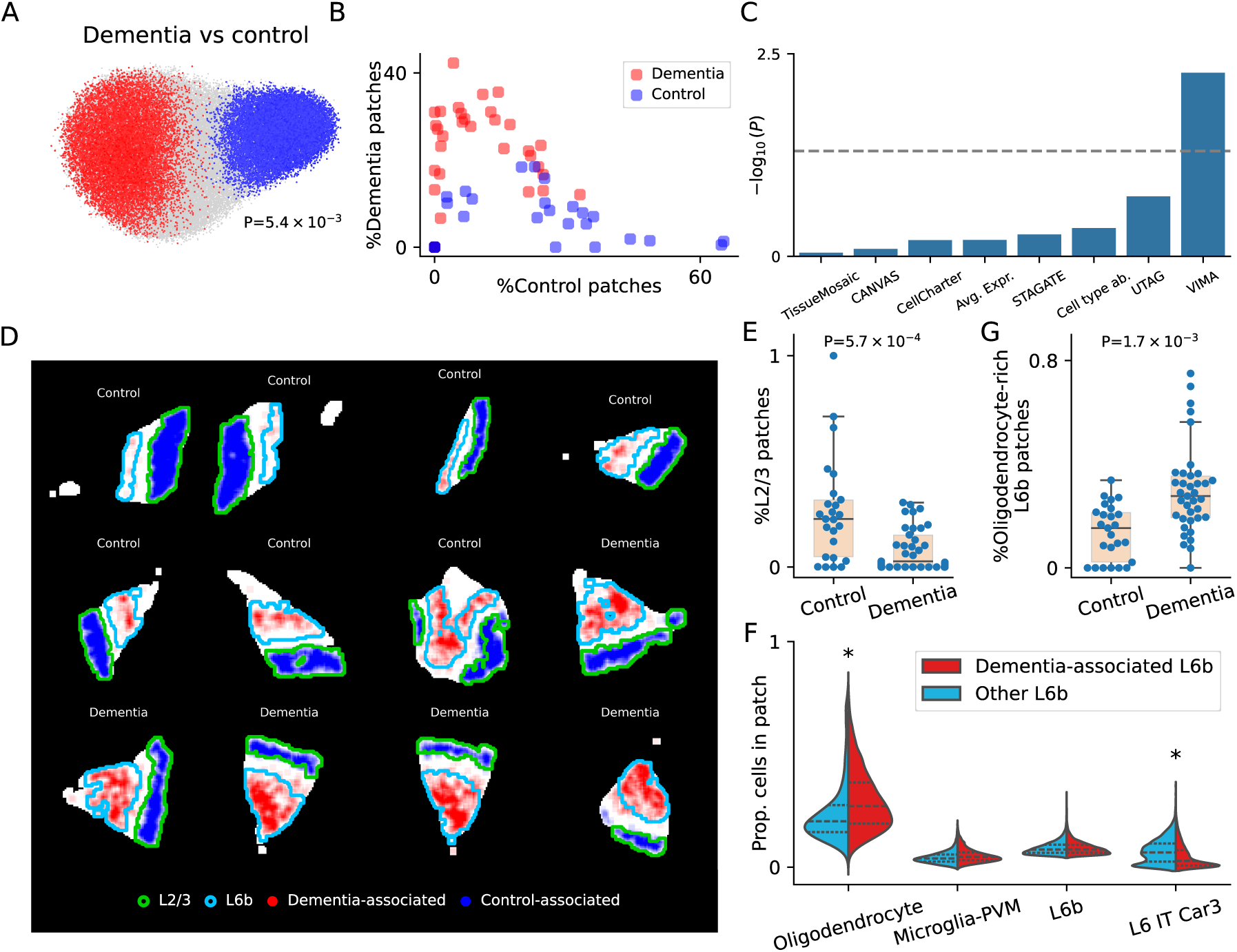
VIMA analysis of Alzheimer’s dataset. A) Results of VIMA case-control analysis for dementia versus control: a UMAP of all the patches in the dataset is generated using the patch fingerprints, and the anchor patches for microniches passing the 10% FDR threshold are colored in proportion to the correlation between the abundance of each microniche and dementia status, with red denoting positive correlations and blue denoting negative correlations. The global P-value for the association test is overlaid. B) Comparison of fraction of dementia-associated vs control-associated patches in each sample. For each sample, the percent of patches in that sample significantly positively associated with dementia is plotted against the percent of patches in that sample significantly negatively associated with dementia. Each sample is then colored by its dementia status. C) Comparison of VIMA’s global P-value for the dementia vs control association to those obtained with tissue representations produced by other methods (Methods). D) Representation of a selected subset of samples with green outlining cortical layer 2/3, light blue outlining cortical layer 6b, red intensity denoting the degree of positive association to dementia for each patch, and blue intensity denoting the degree of negative association to dementia for each patch. E) Distribution of fraction of layer 2/3 patches in dementia vs control samples, with one-sided Kolmogorov-Smirnov P-value overlaid. F) Distributions of cell type proportions of selected cell types among dementia-associated layer 6b patches and non-dementia-associated layer 6b patches. * denotes statistical significance via sample-level permutation test after correction for number of cell types tested (Methods and Supplementary Table 3). G) Distribution of fraction of oligodendrocyte-rich layer 6b patches in dementia vs control samples, with one-sided Kolmogorov-Smirnov P-value overlaid.

The associated patches have clear relationships to cortical architecture and contain both established and novel biological insights. The control-associated patches co-localize very well with cortical layers 2 and 3 (**Fig. 5D**), suggesting thinning of these layers in dementia. This result agrees with multiple existing lines of evidence: a non-spatial single-cell analysis of a larger (N=84) superset of this cohort showed a negative association between abundance of neurons from cortical layers 2/3 and dementia status^47^; prior histology-based^48^ and omics-based studies^47,49^ have found concordant results; and neuroimaging studies have also shown a negative correlation between the thickness of layers 2/3 and measures of cognitive ability^50^. We additionally confirmed this finding in this dataset by manually annotating each patch with a binary label denoting whether it has a high abundance of layer 2/3 neurons (**Methods**); counting the fraction of patches in each sample meeting this criterion and stratifying by dementia status again showed a depletion of layer 2/3 patches in dementia cases (P=5.7e-4 by one-sided Kolmogorov-Smirnov test, **Fig. 5E**). This finding therefore validates VIMA’s ability to recover existing biology in this dataset.

The dementia-associated patches identified by VIMA, in contrast, are novel. These patches lie within the deepest cortical layer – layer 6b – but they do not occupy it uniformly (**Fig. 5D**). To answer the question of what distinguishes the dementia-associated layer 6b patches from other layer 6b patches, we looked for cell type abundance differences between the two sets of patches. We found that the dementia-associated patches contain substantially more oligodendrocytes than the other layer 6b patches (1.33x enrichment, P<1e-4 by donor-level permutation test) and fewer L6 IT Car3 neurons (0.40x depletion, P<1e-4 by donor-level permutation test; **Fig. 5F** and **Supplementary Table 4**). To ensure that these differences were not driven by VIMA simply identifying patches that are more versus less typical of layer 6b, we verified that the dementia-associated patches do not contain different proportions of L6b neurons themselves (**Fig. 5F** and **Supplementary Table 4**). To further confirm this finding, we manually annotating each patch with a binary label denoting whether it has a high abundance of layer 6b neurons and a high abundance of oligodendrocytes (**Methods**); this manual approach reproduced the enrichment (P=1.7e-3 by one-sided Kolmogorov-Smirnov test, **Fig. 5G**). Therefore, our results suggest the existence of a novel oligodendrocyte-rich subset of layer 6b that is more abundant in dementia cases than controls.

Oligodendrocytes have recently been shown in multiple single-cell studies to produce amyloid-β in the cortex^51,52^ and have therefore been implicated in dementia pathogenesis^53^. However, there is also literature about a protective role for oligodendrocytes in this disease^53^. Our finding that dementia-associated patches are enriched for oligodendrocytes specifically within layer 6b raises the possibility that the question of which oligodendrocytes are protective versus deleterious may be partially resolved by accounting for spatial heterogeneity: perhaps the deleterious, amyloidogenic role of oligodendrocytes localizes to layer 6b, which is adjacent to the white matter. This localized, layer-specific enrichment is plausible considering that myelin dysfunction drives amyloid-β deposition in models of Alzheimer’s disease^54^. In this context, our findings therefore suggest ways to advance our understanding of the role of oligodendrocytes in dementia and underscore the importance of spatially resolved studies of cortical architecture in this disease.

### Ablations and assessment of sample-level integration

We conducted ablation studies to determine whether the key components of VIMA are essential. To do this, we constructed 3 VIMA-like methods with progressively fewer of VIMA’s features. We created 1) a version of VIMA that does not have access to sample ID during neural network training and therefore is not able to remove sample-specific bias from the learned fingerprints (“ResNet AE, no cVAE”), 2) a version that is further limited by using a simple two-layer convolutional neural network for its encoder and decoder rather than the ResNet-based architecture used by VIMA (“2-layer ConvNet, Microniches”), and 3) a version that is even further simplified to perform a cluster-based analysis rather than a microniche-based analysis (“2-layer ConvNet, Clustering”). We ran each of these methods on all four of the phenotypes analyzed above (RA subtype, UC vs healthy, TNFi exposure, and dementia), and observed a progressive deterioration of performance as we removed more of VIMA’s central features (**Fig. 6A**). For the RA and UC signals, which are coarser, we find that the most important features are a sophisticated neural network architecture and a microniche-based analysis. In the TNFi and dementia signals, which are more subtle, there is an additional advantage to removal of sample-specific bias, which presumably boosts power by allowing VIMA to produce better integrated representations of tissue patches.

**Figure 6:**
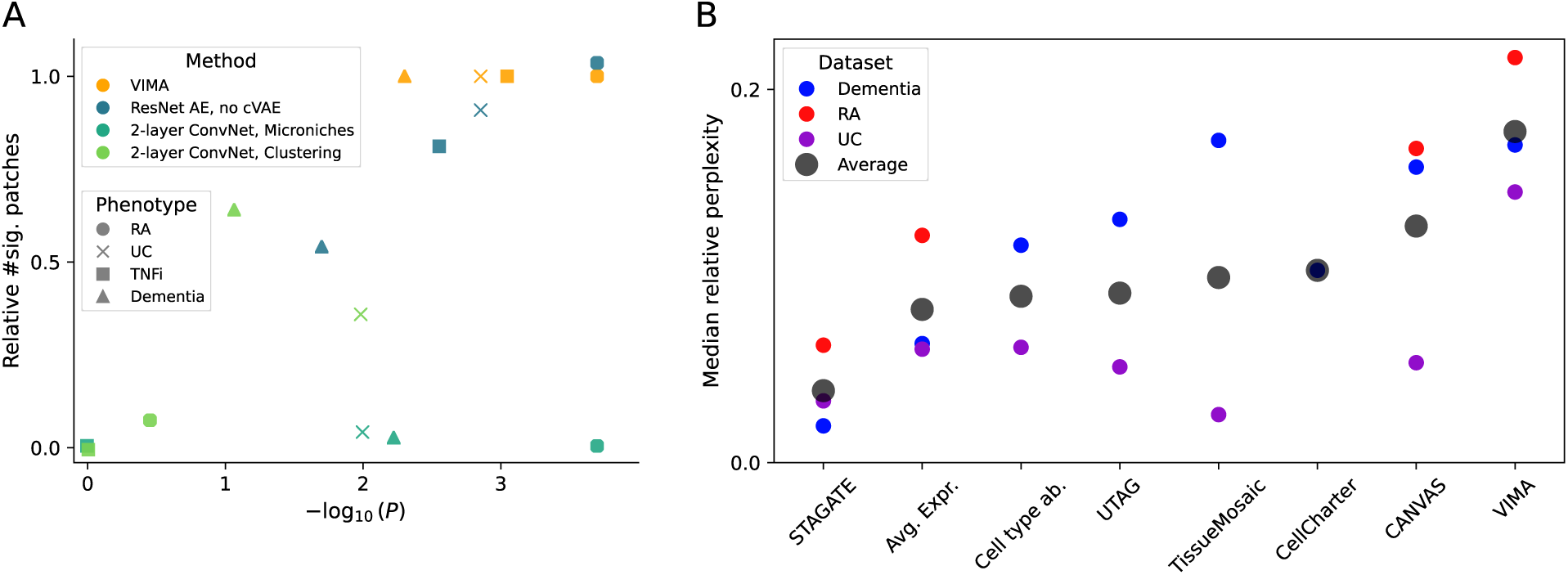
Ablation study and assessment of sample integration on real data. A) Each marker represents either VIMA or one of the three progressively ablated versions of VIMA applied to one of the four phenotypes analyzed in the paper. For each (method, phenotype) pair we plot global P-value for association (x-axis) against the number of patches passing FDR 10% significance relative to standard VIMA. B) For each of the methods we benchmarked and each dataset to which we applied that method, we quantified the degree of integration of the resulting embedding by computing, for each spatial unit in a dataset, the perplexity of the distribution induced across samples by that unit’s neighbors in the relevant nearest neighbor graph, normalizing by the perplexity of all the spatial units, and computing the median across all spatial units in the dataset (Methods). For each method, we additionally plot the average across all the datasets to which it was applied.

In order to verify that both rich patch-level representations and rich sample-level representations improve association testing power, we conducted an additional ablation study using a synthetic phenotype spiked into a downsampled version of the dementia dataset. Here, for ease of simulation, we built a simplified VIMA-like method consisting of a version of VIMA that uses only a single representation at a time (rather than an ensemble of 10). To test the utility of rich patch-level representations, we gave this method progressively less rich patch representations ranging from a 2-layer convolutional neural network to simple patchwide average expression. As the richness of the patch-level representations decreased, we observed a concomitant decrease in power for both the global and local tests, as well as a decrease in spatial accuracy at recovering the true underlying signal (**Supplementary Fig. 8** and **Supplementary Fig. 9**). To test the effect of rich sample-level representations, we then modified each of these methods to use a cluster-based rather than a microniche-based sample representation and again observed decreases in power and spatial accuracy (**Supplementary Fig. 10**).

Assessing the importance of sample-level integration is more difficult in simulation because it is unclear how to simulate realistic technical artifacts. Instead, we conducted an overall assessment, across all three datasets and all the methods we benchmarked, of the degree of sample-level integration achieved by the embeddings generated by VIMA as well as each of the spatial methods we benchmarked. We found that the degree of sample integration achieved by VIMA’s cVAEs was the highest on average across the three datasets, followed by CANVAS, CellCharter, and TissueMosaic (**Fig. 6B**). Correspondingly, when we reduced the incentive of the VIMA-style autoencoder in our simulation framework to integrate samples by reducing the strength of the variational penalty, we again observed a reduction in power and spatial accuracy (**Supplementary Fig. 11**). Taken together, these results suggest that well-powered case-control analysis indeed requires 1) rich patch-level embeddings such as those produced by VIMA’s ensemble of ResNet-style neural networks, 2) a high degree of sample integration, and 3) rich sample-level representations, such as those produced by VIMA’s microniches, that go beyond the information contained in discrete clusters.

## Discussion

Here we introduced VIMA, a method for powerful and accurate statistical case-control analysis of spatial molecular data that can be applied to a broad range of modalities from immunofluorescence microscopy to many-marker protein modalities and spatial transcriptomics. VIMA uses an ensemble of conditional variational autoencoders to extract numerical fingerprints for each patch of tissue in a dataset that quantitatively summarize the biology of that patch. It then uses the fingerprints to define many small, non-discrete, and data-dependent groups of highly similar patches called microniches, and asks whether the abundance of these microniches correlates with case-control status. The end result is 1) a tensor of samples by microniches by autoencoders called the microniche abundance tensor (MAT) that summarizes the abundance patterns of different microniches across the samples, 2) a global P-value for association to case-control status, and 3) a subset of the patches that explain that association at a specified false discovery rate, with a directional effect size for each patch. VIMA does this while accounting for sample-specific artifacts, allowing for inclusion of covariates, requiring minimal parameter tuning, and running efficiently even on datasets with 50-100 samples and hundreds of millions of transcripts. VIMA is different from the deep learning-based methods that have become popular in digital pathology^55,56^, which typically are trained on thousands or even millions of patient samples and focus on patient-level prediction of disease status in a clinical setting rather than statistically defining the specific spatial structures that contribute to that disease in a much smaller research cohort.

In the course of developing VIMA, we observed that individual cVAEs trained on the same data could produce representations that differed due to the stochasticity of neural network training. This suggested that each cVAE might capture a distinct and informative perspective on the data. By training an ensemble of ten cVAEs and analyzing them jointly, VIMA both leverages these multiple complementary views of the dataset to improve detection of biologically meaningful signal and also increases replicability.

We applied VIMA in three datasets, each from a different spatial modality. In each case, VIMA detected novel biological signals, including: previously unknown spatial heterogeneity within rheumatoid arthritis synovium alongside new spatial characterization of known rheumatoid arthritis disease subtypes, a statistically robust signature of TNF inhibition in ulcerative colitis characterized by replacement of lymphoid aggregates with disorganized mesenchymal activity, and a dementia-associated tissue niche consisting of oligodendrocyte-rich areas in cortical layer 6b. In each case, we showed using detailed benchmarking that these signals are not detectable using the tissue representations generated by current state-of-the-art methods. **Supplementary Fig. 12** summarizes these results across all four case-control signals tested, demonstrating both VIMA’s power as well as its ability to work across different spatial technologies. We conducted ablation studies demonstrating that each of VIMA’s key ingredients – a rich patch representation based on neural network architectures from image processing, an ability to avoid overfitting to sample-specific idiosyncrasies, and a microniche-based sample representation that allows for granular case-control analysis — is necessary for its superior performance.

In our benchmarking study, we used each tissue representation in the standard fashion by clustering spatial units according to their representations then testing each cluster for association and correcting for multiple testing (**Methods**). Variations on this strategy, such as performing the association testing across clusters using a single global test statistic akin to VIMA’s global test, harmonizing the representations across samples prior to clustering, or feeding the representations into a VIMA-like microniche framework led only to modest improvements in performance, with VIMA still retaining a superior ability to detect these signals (**Supplementary Fig. 13**, **Supplementary Fig. 14**, and **Supplementary Fig. 15**). Thus, both the way in which VIMA creates patch fingerprints and the way that it uses those fingerprints are important for its performance. We emphasize, however, that the majority of the methods we benchmarked were not developed explicitly for statistical case-control analysis but rather for tissue annotation, which they have already been shown to do extremely well. Our benchmarking has no bearing on the biological accuracy of these annotations in any one sample, which we believe is high; rather, it is meant to assess the degree to which such annotations can be easily repurposed for large-scale statistical case-control analysis.

VIMA treats spatial data directly as a set of images and does not rely on segmentation into cells, cell types, and tissue regions. This has the advantage of easily accommodating a broad range of spatial modalities, avoiding the need for error-prone discretization that may not accurately model the data, and reducing the number of user-dependent parameter choices (e.g., number of cell types, tolerance of cell segmentation, etc.). It also allows for simple integration of modalities if more than one is available: each modality’s marker/gene channels can simply be concatenated with the others. As our understanding of this approach improves, it may become possible to also utilize pre-trained models, such as the foundation models that exist today for processing hematoxylin and eosin (H&E) data^55,56^, to boost performance further.

VIMA has several limitations. First, although VIMA runs relatively quickly, this speed requires a GPU and using VIMA without one can be time-consuming; this can be ameliorated by using multiple CPUs, but obtaining enough CPUs to match one GPU may be possible only on a computing cluster. Second, although VIMA’s performance appears robust to choices of many of its parameters, we do feel that dramatically different choices of patch size (e.g., 40um vs 400um in length) could lead to emphasis of different types of signals. Third, since VIMA’s output is not a set of defined and manually annotated tissue niches, understanding the driving biology behind a VIMA result requires thought; we maintain, however, that as with other genomic modalities this process is always necessary to produce biological insight from spatial data, whether before or after the case-control analysis is performed. Applying this careful thought after the spatial regions of interest are delineated may be advantageous. Finally, while VIMA’s relatively unconstrained approach allows it to detect a broad range of case-control signals, more targeted methods tailored to specific case-control signal types may outperform VIMA when their modeling assumptions align closely with the underlying ground truth in the data.

Despite these limitations, VIMA is a sensitive and accurate way to identify disease-relevant spatial features across a wide range of spatial modalities that is unique in leveraging the power of deep learning within a rigorous and flexible statistical framework. As spatial molecular datasets grow in sample size and dimensionality, methods that can give a detailed picture of variation at the sample level will become increasingly important for translating these unprecedented datasets into advances in our understanding of disease.

## Methods

### Variational Inference-based Microniche Analysis

#### Intuition

Variational inference-based microniche analysis is built on two technical concepts: 1) the patch fingerprint, which is a numerical representation of a tissue patch learned by a variational autoencoder that can be used to quantitatively assess the similarity of pairs of patches, and 2) the microniche, which is a small set of tissue patches from multiple samples that are biologically very similar to each other. Broadly, our method proceeds by using the patch fingerprints to construct the microniches and then finding case-control differences by searching for microniches whose abundance correlates with case status in a statistical hypothesis testing framework.

In the remainder of this section, we establish notation and then proceed to provide a detailed description of 1) how VIMA pre-processes the data, with attention to minimizing pixel-level batch effects, 2) how VIMA extracts patch fingerprints using conditional variational autoencoders while accounting for spatial batch effects, 3) how VIMA defines one set of microniches for each of the ten autoencoders it trains, and 4) how VIMA conducts statistical association testing on the microniches, including meta-analyzing over the ten autoencoders.

#### Notation and assumptions

Let {*S*_1_, …,*S_N_*} be a set of *N* spatial molecular samples with *M* different markers or genes measured for each sample. We assume that each sample has been rasterized into pixels of size *r* x *r*, and we represent each sample *S_n_* as a tensor of shape (*X_n_*, *Y_n_*, *M*) whose (*x*, *y*, *m*)-th entry is either the intensity of marker *m* in the *x,y*-th pixel (for non-transcript-based modalities) or the total count of gene *m* in the *x,y*-th pixel (for transcript-based modalities). For ease of exposition, we occasionally refer below to this number as intensity even for transcript-based modalities. In all analyses in this paper, we set *r* to be 10 micrometers; however, smaller values are in principle possible and could reveal aspects of tissue architecture that play out on smaller length scales.

#### Pre-processing and pixel-level batch correction

VIMA pre-processes data with the goals of a) differentiating tissue from background, b) reducing a potentially large number of markers into a more manageable set of “meta-markers”, each of which is a linear combination of the original markers, and c) removing unwanted sample- or batch-specific artifacts from the meta-markers.

To accomplish (a), a dataset-dependent number is computed for each pixel summarizing the total amount of signal in that pixel: this may be the total number of transcripts in that pixel (for transcript-based modalities), the sum of all marker intensities divided by the sum of the intensities of all negative control stains (for non-transcript-based modalities), etc. VIMA then uses this number to segment each slide into foreground (i.e., tissue) and background. For transcript-based modalities, VIMA simply applies a threshold of 10 transcripts per pixel. For non-transcript-based modalities, in which non-specific background staining is a prominent feature, VIMA instead uses Otsu thresholding^57^. Once the foreground is defined, all background pixel intensities are set to 0 for all markers.

To accomplish (b), VIMA first log-normalizes all non-empty pixels: it computes the median total intensity *q* of all the pixels in the dataset and normalizes each pixel *p* according to the formula

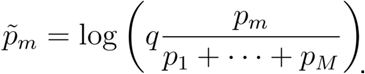

For each pixel, it then averages the intensity values of any non-empty pixels in the 5×5 square centered at the pixel in question, and it performs PCA on the resulting “meta-pixels”. This PCA yields a set of PCs defined across all the markers. VIMA uses these loadings to then convert each of the original normalized pixels from a length-*M* vector of raw markers to a length-*K* vector of meta-markers defined by the loadings of the top *K* PCs. The default value of *K* is 10, but when a dataset contains fewer than 10 markers, we use *K*=5 instead.

To accomplish (c), VIMA removes sample-specific and batch-specific artifacts by applying the Harmony algorithm^34^ with default parameters to the meta-marker representation of all the non-empty pixels in the dataset.

#### Featurization of patches using multiple conditional variational autoencoders

Once the pixels are pre-processed, VIMA converts each sample into a set of tissue patches of size 40×40 pixels. (Depending on cellular density, each tissue patch typically contains tens to hundreds of cells. Since the tissue patches overlap, a 1×1cm tissue fragment would yield 10,000 tissue patches with meta-marker information on 1 million pixels.) All patches for which fewer than 20% of the pixels contain tissue (as opposed to background) are then discarded.

VIMA then trains ten conditional variational autoencoders in parallel to each learn a *C*-dimensional latent representation from which it can reconstruct the tissue patches, where *C*=100 by default. The loss function that is optimized is:

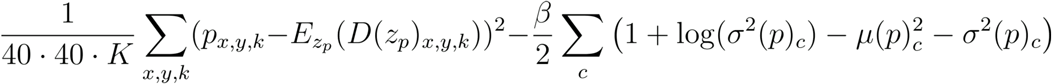

where *p* is the original patch (represented as a 40×40x*K* tensor), *D* represents the decoder, *μ*(*p*) and *σ*^2^ (*p*) are the *C*-dimensional mean and variance respectively of the multivariate normal distribution specified by the encoder as the encoding of *p*, 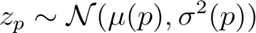 is a sample from that distribution, and *β* determines the relative weights of the two components of the loss function; a small value of *β* prioritizes reconstruction while a large value of *β* prioritizes semantics of the latent space. The loss function is optimized stochastically in the standard way using the reparameterization trick^58^. Because the runtime of the training process has non-trivial contributions both from backpropagation and from serving the training instances, training all ten autoencoders in parallel takes significantly less than 10 times as long as training them in series.

The autoencoders’ architecture is shown in **Supplementary Fig. 16**. This architecture is similar to that of the ResNet^36^ family of neural networks, with the following two modifications. First, it has fewer layers because the patches are of size 40×40 rather than 224×224. Second, the autoencoder is a conditional variational autoencoder^35^, meaning that both the encoder and decoder are given access to sample-level covariates whose influence on the latent representation the user would like to minimize. This is accomplished by allowing the encoder and the decoder to each learn a 4-dimensional embedding *w^n^* for sample *n* that is a function of a one-hot encoding of sample ID together with any other covariates supplied. This is converted into four new spatially homogeneous channels by setting all spatial locations in the *i*-th new channel to 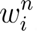. These new channels are then concatenated to the model’s representation at each layer. In this way, the regularization imposed by the variational penalty incentivizes the encoder to leverage the new channels to remove any sample-specific information, and incentivizes the decoder to then supply that information back during its reconstruction. This method of incentivizing integration by giving the encoder and decoder access to sample-level information has been successfully used in single-cell integration^59^.

In simulation, we show that performance does not strongly depend on *β* as long as it’s not extremely small (**Supplementary Fig. 11**). Accordingly, in practice we simply set *β* = 10^−5^ with the recommendation that if the patches are not well integrated across samples then *β* should be increased by one order of magnitude at a time until this is achieved. (We have not had to do this in the datasets analyzed in this paper.)

We train the autoencoders using standard practice, with 80% of the patches used for training and 20% used for validation. We use a batch size of 256 patches, a default learning rate of 10^-3^ with an exponential decay parameter of 0.9 per epoch, KL warmup over the first 5 epochs, and 20 epochs of training with option to extend if the validation loss has not plateaued. These settings were uniform across all datasets processed.

#### Definition of microniches

Once the autoencoders are trained, they are each run on all *P* patches in the dataset whose tissue density (the fraction of pixels containing tissue) is greater than some threshold *α* to produce a *C*-dimensional vector *z_p_* for each patch *p*; *α* is set of 0.5 by default. Each set of embeddings *z_p_* is then used to construct a nearest-neighbor graph of all *P* patches using scanpy’s neighbors function^60^.

The microniches are then defined similarly to how transcriptional neighborhoods are defined in our previous work in non-spatial single-cell analysis^37^. That is, we define one microniche for each patch *p* and each autoencoder *e* using the similarity implied by the structure of the nearest-neighbor graph *G_e_* generated by the patch fingerprints created by *a*: for every other patch *p’*, the degree of belonging to the microniche anchored at patch *p* is the probability that a random walk in *G_e_* starting at patch *p’* will arrive at patch *p* after *s* steps. The transition matrix of the random walk is *A_e_* + *I* where *A_e_* is the adjacency matrix of *G_e_* and *I* is the identity matrix. The number of steps *s* is chosen adaptively by stopping the random walk once enough microniches are no longer gaining representation from new samples with each step of the walk. This is accomplished by tracking the median kurtosis across microniches of the distribution across samples of each microniche, and stopping when the decrease in this number with each step of the random walk falls below a hardcoded threshold.

Applying the above procedure for each sample in the dataset allows us to build the MAT, which formally is an *N* × *P* × 10 tensor whose *n,m,e*-th entry equals the expected fraction of patches from sample *n* that would arrive at patch *p* after *s* steps in *G_e_*, that is, the relative abundance of microniche *p* in sample *n* according to the fingerprints generated by the autoencoder *e*. The MAT can be computed quickly using sparse matrix multiplication since the matrix *A_e_* + *I* is sparse. As with prior work^37^, our software allows for covariates to be projected out of the MAT.

#### Statistical association testing

Given the MAT *Q* and a sample-level attribute of interest *y*, VIMA performs two types of association testing: a *global* association test, which is meant to assess for any aggregate relationship between abundance of the microniches as captured by *Q* and *y*, and a *local* association test, which is meant to find specific microniches whose abundance correlates with *y*.

VIMA begins by standardizing *Q* along its second dimension (e.g., each microniche is standardized). It then computes a matrix *C* of dimension 10 × *P* whose *e,p*-th entry is the correlation across samples between the phenotype *y* and the abundance of each sample in the *p*-th microniche according to autoencoder *e*.

To compute the local test, VIMA then computes, for each patch *p*, a meta-analyzed correlation *ρ_p_* using the ratio of the third to second moments, i.e.,

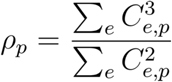

which yields a single length-*P* vector *ρ* with one meta-analyzed correlation per patch. The motivation for this meta-analysis is that we would like to compute the average correlation across all 10 auto-encoders, but we want to allow for the possibility that for a given patch *p*, some autoencoders have learned more useful structure than others. We therefore weight our average by the square of each respective correlation. To assess statistical significance, VIMA then does the same procedure for a large number of random permutations of *y*. It then computes, for each potential value *ρ*\* that might be used as a cutoff for statistical significance, an empirical estimate of the false discovery rate at that threshold. That is, it naively estimates

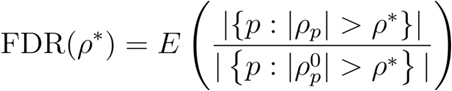

Where 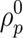 is a random variable that represents the meta-analyzed correlation between the microniches anchored at patch *p* and a random permutation of *y*. To account for the presence of multiple samples from the same donor, which is common in spatial datasets, the permutation scheme used preserves the property that samples from the same donor have the same sample-level attribute.

To perform the global test, VIMA computes, across all patches *p* and autoencoders *e*,

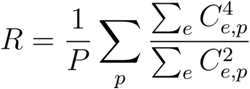

which quantifies the total amount of correlation signal in the matrix *C*. It then computes a large number of null instances of *R* using the same null permutation scheme described above and compares R to these to produce a single global P-value for association.

### Null simulations

To assess the calibration of VIMA, we performed simulations using a real spatial transcriptomics dataset from the SEA-AD dementia cohort^47^. This dataset consists of S=75 postmortem medial temporal gyrus samples from N=27 patients, 15 of whom had dementia. Each sample was profiled using MERFISH, yielding spatial information about M=140 genes at single transcript resolution, with a total of 516 million transcripts profiled. For the simulation in **Supplementary Fig. 1**, we generated 500 simulates in which the samples were left unmodified and a random binary case-control label was assigned to each sample. We then quantified type I error at level *α* = 0.05 by computing the fraction of null simulates in which VIMA rejected the null hypothesis.

### Analyses of real dataq

We analyzed three real datasets: synovial biopsies from patients with RA profiled with immunofluorescence microscopy (S=27 samples from N=22 donors, M=7 markers profiled)^39^; colonic biopsies from patients with and without ulcerative colitis profiled with CODEX (S=42 samples from N=34 donors, M=52 markers profiled)^19^; and post-mortem medial temporal gyrus samples from patients with and without dementia profiled with MERFISH (S=75 samples from N=27 donors, M=140 genes profiled)^47^.

In addition to VIMA, we also analyzed these datasets with a suite of benchmarking methods: CellCharter^22^, STAGATE^26^, UTAG^21^, CANVAS^32^, TissueMosaic^33^, a “patchwide average expression” baseline in which each tissue patch was featurized by the patchwide average of the harmonized meta-markers generated by VIMA’s pre-processing, and a “patchwide cell type abundance” baseline in which each tissue patch was featurized by the relative abundances of each cell type in that tissue patch. These methods were chosen to represent a diversity of approaches to single-modality tissue annotation that are prevalent in the field, including both neural network-based and non-neural network-based methods, and including both methods that work from segmented cells as well as methods that work from raw expression data. Each method was run on each dataset with the following exceptions: in the RA dataset, we were unable to apply CellCharter, UTAG, TissueMosaic, and the patchwide cell type abundance baseline because we did not have access to segmented cells; and in the UC dataset, we were unable to apply CellCharter because its pre-processing step for CODEX data, which integrates expression profiles of all cells using trVAE^61^, exceeded our runtime limit of 3 days.

#### Analysis of rheumatoid arthritis dataset

We obtained the RA dataset from the AMP RA/SLE consortium. We pre-processed the data with the default preprocessing workflow described above, using the ratio between the summed intensity of all 7 markers and the intensity of the auto-fluorescence channel to summarize the total amount of tissue signal in each pixel. After log-normalization, PCA across meta-pixels, and harmony as described above, this resulted in 5 (harmonized) meta-markers per pixel, and a mask for each sample that defines which pixels are assumed to come from tissue. We then ran VIMA on the dataset with default settings, except that we set *α* (the fraction of pixels in a patch that must be non-empty for VIMA to include it in the case-control analysis) to 0.8 due to the difficulty accurately distinguishing tissue from nonspecific background staining in this dataset as well as the highly irregular borders of many of the samples.

In addition to using VIMA, we also generated tissue annotations for this dataset using CANVAS^33^, STAGATE^26^, and our patchwide average expression baseline. (The additional methods in our benchmarking suite all required cell segmentation, which was not available for this dataset; see above.) For details of how these methods were applied, see “Details of benchmarking” below.

To compute the degree of sample integration in each method’s embedding for **Fig. 2B**, we constructed a nearest neighbor graph on the spatial units used by each method using whatever embedding that method assigned to those spatial units and using scanpy’s default parameters. For each method, we then quantified, for each spatial unit, the perplexity of the distribution over samples induced by that unit’s neighbors in the graph, and divided this by the perplexity of the distribution over samples induced by all the spatial units in the graph; we refer to this as “relative perplexity across samples”.

To apply principal components analysis to VIMA’s MAT for **Fig. 3A**, we converted the MAT from a *N* = *P* = 10 tensor into an *N* × 10*P* matrix and applied standard PCA to the matrix. (We note that in the future tensor decomposition could be used instead, which would give directions of variance explained by multiple autoencoders; we did not pursue this here.)

For the case-control analysis, we used each sample’s CTAP labels, determined using separate non-spatial single-cell RNAseq. The 6 CTAPs are T, TB, TM, TF, EFM, and F and reflect preponderances of various combinations of T cells (T), B cells (B), fibroblasts (F), and myeloid cells (M). We defined a sample to be “stromal” if it is in the union of all the fibroblast-containing CTAPs, i.e.,

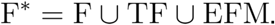

and “immune” otherwise. This was the phenotype that we used as input to VIMA’s case-control analysis, resulting in a global P-value for association to stromal status and a set of microniches significantly associated (both positively and negatively) with stromal status at FDR 10%.

To produce the benchmarking results in **Fig. 3C**, we ran an analogous case-control analysis using the tissue annotations produced by each of the other three comparator methods used in this dataset. For each method, we clustered the spatial units used by that method according to the embedding that method assigned to those spatial units. If a method provided its own clustering methodology we used that methodology; otherwise we used leiden clustering with default parameters. We then computed, for each cluster, the squared correlation between abundance of that cluster in each sample and the phenotype. We then produced a P-value for each cluster using null permutations and obtained a global P-value by applying a Bonferroni correction across clusters. We also tried an alternative approach of performing the association testing across clusters using the sum of squared correlations across the clusters as a global test statistic and comparing to a permutation-based null distribution, akin to VIMA’s global test. This yielded only slightly better results (**Supplementary Fig. 13**). We additionally tried harmonizing each method’s embedding across samples prior to association testing (**Supplementary Fig. 14**) and also feeding each method’s embedding directly into VIMA to yield a microniche-based association test (**Supplementary Fig. 15**).

To assess which markers have higher average intensity in VIMA’s stromal-associated patches, we computed the average intensity of each marker in each patch. We then estimated the distribution of each marker across all the patches identified by VIMA as typical of the stromal samples and across all the patches identified by VIMA as typical of the immune samples (**Supplementary Fig. 2**).

We next clustered the positively and negatively associated patches, respectively, into 2 clusters each. We compared average marker levels in the same way that we did for the stromal-associated patches (**Supplementary Fig. 3** and **Supplementary Fig. 4**). We then computed the enrichment of each CTAP in each cluster as follows. For each of the CTAPs *y* with more than 50 patches represented among, e.g., the stromal-associated patches, and for each of the clusters *c*, we calculated the number of stromal-associated patches in cluster *c* from all samples with CTAP *y*. We then computed the log-odds-ratio, for each CTAP, of a patch from that CTAP belonging to cluster *c* versus not belonging to cluster *c*. We calculated standard errors for the log-odds-ratio using the formula for the asymptotic variance formula for log-odds-ratios computed from contingency tables (1/*a* + 1/*b* + 1/*c* + 1/*d* where *a*, *b*, *c*, *d* are the four counts used). For the immune-associated patches, there were not enough patches from stromal CTAPs to perform the enrichment analysis for the stromal CTAPs. For the stromal-associated patches, we were able to perform the enrichment analysis for all CTAPs, but only the immune CTAPs had appreciable enrichments. The full enrichment results for the stromal-associated patches are presented in **Supplementary Fig. 5**.

#### Analysis of ulcerative colitis dataset

We obtained the UC dataset in tiff format from the authors of the original study^19^. In addition to spatial data, each patient also has metadata specifying whether they have UC or not, whether they were on a TNF inhibitor at the time of their biopsy, and whether they were on a TNF inhibitor in the past. In addition to the 52 CODEX markers, each sample also had 21 channels consisting of Hoechst stains that were used to register the markers across different rounds of imaging in the CODEX protocol. We noticed that these stains induced substantial inter-sample batch effects so we removed them prior to normalizing the pixel intensities in the dataset.

We pre-processed the data with the default preprocessing workflow described above, using the summed intensity of all the CODEX markers and the nuclear stains to summarize the total amount of tissue signal in each pixel, but using only the CODEX markers for the dimensionality reduction from markers to meta-markers. This resulted in 10 (harmonized) meta-markers per pixel, and a mask for each sample that defines which pixels are assumed to come from tissue. We then ran VIMA on the dataset with default settings to check for association to UC vs healthy status, removing from the case-control analysis (but not from the training of the autoencoder) any sample that either was on a TNF inhibitor or had been on one in the past in order to maximize the contrast between ulcerative colitis and healthy tissue. This resulted in a global P-value for association as well as a set of microniches significantly associated (both positively and negatively) with UC status at FDR 10%.

In addition to using VIMA, we also generated tissue annotations for this dataset using CANVAS^32^, STAGATE^26^, UTAG^21^, TissueMosaic^33^, our patchwide average expression baseline, qqand our patchwide cell type abundance baseline. (CellCharter^22^, whose proteomics workflow uses trVAE^61^ to integrate the expression profiles of the cells, did not finish within our runtime limit of 3 days due to the trVAE step.) For details of how these methods were applied, see “Details of benchmarking” below.

To assess which markers have higher average intensity in the UC-associated patches, we computed the average intensity of each marker in each patch. We then estimated the distribution of each marker across all the UC-associated patches identified by VIMA as well as the healthy-associated patches.

To identify patches typical of treatment with TNF inhibition, we took only the patches found by VIMA to have a significant positive association with UC and re-built new nearest neighbor graphs using the patch fingerprints of these patches. We then used these new nearest-neighbor graphs to define a new MAT and ran our association testing with this new MAT with a phenotype of “chronic/prior TNF inhibition” defined as presence of past use of TNF inhibition. qWe then repeated the same workflow described above to identify markers typical of the positively and negatively associated patches.

To show that the chronic/priorTNFi-vs-never/newTNFi signature was distinct from the UC-vs-healthy signature, we selected all the patches found to be significantly chronic/priorTNFi-associated and compared their UC microniche coefficients to those of the patches that were significantly never/newTNFi-associated (**Supplementary Fig. 6**). If the two signatures were similar, we might expect the chronic/priorTNFi-associated microniches to have lower correlations to UC on average than the never/newTNFi-associated patches; instead, we saw nearly identical distributions, with the median UC correlation actually being slightly higher among the chronic/priorTNFi-associated microniches.

#### Analysis of dementia dataset

We downloaded the Alzheimer’s dataset from the SEA-AD website. In addition to spatial data, each patient was also labeled as either having dementia or not. To enable comparison to the local averaging paradigm as instantiated through local cell-type abundances, we performed cell segmentation on the data as described in ref.^62^ and then annotated those cells using the Allen Institute’s MapMyCells tool (https://portal.brain-map.org/atlases-and-data/bkp/mapmycells) qusing the matching “10x Human MTG SEA-AD (CCN20230505)” reference taxonomy and the “SEA-AD Correlation Mapping” algorithm. As a result, each cell was annotated as having a spatial location and a cell type at several specific cell-type resolutions.

We pre-processed the data with the default preprocessing workflow described above, using the total transcript count per pixel to summarize the total amount of tissue signal in each pixel. This resulted in 10 (harmonized) meta-markers per pixel, and a mask for each sample that defines which pixels are assumed to come from tissue. We then ran VIMA on the dataset with default settings to check for association to dementia vs non-dementia status, yielding a global P-value for association and a set of significant microniches at FDR 5%.

In addition to using VIMA, we also generated tissue annotations for this dataset using CANVAS^32^, STAGATE^26^, UTAG^21^, TissueMosaic^33^, CellCharter^22^, our patchwide average expression baseline, and our patchwide cell type abundance baseline. For details of how these methods were applied, see “Details of benchmarking” below.

To determine whether the positively and negatively associated microniches coincide with specific cortical layers, we computed for each patch the proportion of cells in that patch belonging to each cell type. We used this information to spatially delineate as Layer 6b all patches for which “L6b” comprised at least 5% of the cells in the patch and to delineate as Layer 2/3 all patches for which “L2/3 IT” comprised at least 30% of the cells in the patch.

To determine which cell types were most enriched in the dementia-associated patches compared to other layer 6b patches, we computed the proportion of cells in each patch coming from each of the 24 annotated cell types. We then compared the median cell type abundance for each cell type between the dementia-associated patches and the other layer 6b patches. We tested these differences for significance by randomly flipping, for each sample, the association status (dementia-associated vs not) of all of the layer 6b patches in that sample and using this to compute a null distribution for our statistic. This yielded the P-values in **Supplementary Table 4**.

To confirm our interpretations of the dementia- and non-dementia-associated patches, we used the cell type proportions of each patch to define two types of patches: layer 2/3 patches, defined as those patches with more than 30% of cells belonging to the “L2/3 IT” cell type, and oligodendrocyte-rich layer 6b patches, defined as those patches with more than 5% of cells belonging to the “L6b” cell type and more than 10% of cells belonging to the “Oligodendrocyte” cell type. We then computed the proportion of patches from each of these two categories in each sample and compared the distributions among cases and controls using a one-sided Kolmogorov-Smirnov test.

To compare to a non-spatial analysis of this same dataset, we represented each sample using a vector containing the proportion of cells in that sample belonging to each of the 24 annotated cell types and used correlation to case/control status with permutation-based empirical false discovery rates computed analogously to those of VIMA’s (**Supplementary Fig. 7**).

### Benchmarking, ablations, and simulations

#### Details of benchmarking

We provide below details of how each method in our benchmarking suite was run, followed by details of how each method’s embedding was used for case-control analysis.

##### STAGATE

Since STAGATE’s spatial unit is a visium-sized tissue region for which it is supplied with an average expression profile, we generated 100×100um patches for each dataset and computed the average expression profile of each patch. We then ran STAGATE with default parameters, rad_cutoff=150um, and n_epochs=1000.

##### CANVAS

We ran CANVAS’s preprocessing code directly on the 10um-resolution pixelated counts matrices generated by VIMA’s rasterization procedure, specifying the average of all the channels as the “background” channel per the documentation. We then trained the CANVAS model, stopping training at 200 epochs due to runtime.

##### Patchwide average expression

For each VIMA patch, we computed the average of the harmonized meta-markers within that patch.

##### UTAG

We obtained segmented cells with cell-level expression data from the authors of the UC study and from the online portal associated with the Alzheimer’s disease study, and we ran UTAG on these cells according to the “batch mode” tutorial, setting max_dist=15, and normalization_mode=’l1_norm’ per that tutorial.

##### CellCharter

We obtained segmented cells with cell-level expression data from the authors of the UC study and from the online portal associated with the Alzheimer’s disease study, and we can CellCharter on these cells with default parameters and following the online tutorials associated with the method, which recommended integration of the cells with scVI^63^ for the spatial transcriptomic data and trVAE^61^ for the CODEX data. However, trVAE did not finish training on the UC dataset with default parameters in 3 days, and so we did not include CellCharter results for that dataset.

##### TissueMosaic

We obtained cell types from the authors of the UC study for the segmented cells in that study, and we used the MapMyCells tool as above to call cell types for the segmented cells in the Alzheimer’s disease study. We then gave these as input to TissueMosaic and trained the model with default parameters except for modest adjustments to the training schedule due to runtime considerations (num_workers=16, param_momentum_epochs_end=700, warm_up_epochs=70, warm_down_epochs=70, and max_epochs=700) and global_size=100 rather than 96 (after discussion with the authors, in order to match the TissueMosaic patch size to VIMA’s patch size).

##### Patchwide cell type abundances

We obtained cell types from the authors of the UC study for the segmented cells in that study, and we used the MapMyCells tool as above to call cell types for the segmented cells in the Alzheimer’s disease study. We then computed the relative abundance of each cell type in each of VIMA’s patches.

To evaluate each method’s embedding, we clustered the spatial units generated by each method either using Leiden clustering with default parameters or, if available, using whatever clustering was performed by the method in question. We then produced an *N x K* matrix, where *N* is the number of samples and *K* is the number of clusters, whose *n,k*-th entry is the relative abundance of cluster *k* in sample *n*. We then computed the correlation *C_k_* between the *k*-th cluster the phenotype for each *k*. We produced one P-value per cluster a by comparing *C_k_* to null distribution obtained by permuting the sample labels while preserving identical labels for samples from the same donor. We then computed a global P-value for association by applying a Bonferroni correction to the set of resulting P-values. We computed empirical false discovery rates for each cluster in the same way that VIMA does for microniches, and used this to quantify how many spatial units passed 10%-FDR significance.

In addition to the above approach, which was used for all main text figures and whose results are summarized in **Supplementary Fig. 12**, we separately implemented the following alternative approaches: 1) We produced a global P-value for association in the same way that VIMA does by summing the squares of the *C_k_* and comparing to the permutation-based null distribution. 2) We ran Harmony^34^ on the embeddings generated by each method and then performed the main cluster-based analysis. 3) We generated microniches from the embeddings produced by each method and then fed these into VIMA, leading to a global P-value and local FDRs. These analyses are summarized in **Supplementary Fig. 13**, **Supplementary Fig. 14**, and **Supplementary Fig. 15**.

#### Ablation study

For each dataset, we trained the following ablated models:

1. VIMA_nosid, a method identical to VIMA except in that the autoencoder does not have access to sample ID.
2. VIMA_nosid_noresnet, a method identical to VIMA_nosid except that the autoencoder has as its encoder and decoder a simple 2-layer convolutional neural network with 256 and then 512 colors in its intermediate representations and 100 latent dimensions in its bottleneck.
3. VIMA_nosid_noresnet_nomicroniches, a method identical to VIMA_nosid_noresnet except that instead of representing each sample using the microniche abundance tensor, each sample was represented using clusters generated via the fingerprints produced by a single autoencoder.

For each model and each dataset, we reported global P-values and number of significant patches at FDR 10%; for some models, the TNFi association was not runnable because there qwere not enough UC-associated patches to perform the analysis; in this case, we report a P-value of 1 and 0 significant patches.

#### Comparison of integration across methods

For each of the methods profiled for each dataset, and for each spatial unit, we quantified sample integration using relative perplexity as described in the rheumatoid arthritis dataset. We then examined the distribution of relative perplexities for each (method, dataset) pair and plotted these in **Fig. 6B**.

#### Simulations to assess richness of patch- and sample representations

We performed our simulations using the Alzheimer’s dataset, though for computational convenience, we downsampled the dataset to one sample per patient for the simulation study.

We simulated three signals: A) a scenario in which cases have focal cellular aggregates and controls do not, B) a scenario in which cases have focal cellular aggregates while controls have a diffuse infiltrate, C) a scenario in which cases have focal cell aggregates while controls have a striated tissue structure.

To simulate these signals, we first defined a region in each sample corresponding to cortical layer 2/3 by using the annotated cell locations published with the dataset: we identified the pixel location of all layer 2/3 neurons and used standard imaging transforms (closing with radius 40 pixels followed by opening with radius 10 pixels) to define a contiguous region around areas with high density of such pixels. We then identified all pixels lying within the layer 2/3 region annotation whose second meta-marker was above 5 and defined these as *foci* around which we would spike in any new signals. The spiked in signals were: for (a) addition to meta-marker 2 of a Gaussian blur of diameter 7 pixels around each focus in case samples; for (b), addition to meta-marker 2 of a Gaussian blur of diameter 7 pixels around each focus in case samples and a Gaussian blur of diameter 51 pixels around each focus in control samples; for (c), addition to meta-marker 2 of a circle of diameter 7 pixels around each focus in case samples and a rectangle with shape 1×33 pixels around each focus in control samples.

For each non-null signal type, we added increasing amounts of noise by randomly re-assigning the case/control label of a given fraction *h* of the samples for *h* ∈ {0, 0.1, …, 0.5} We performed 50 simulates for each signal type at each noise level.

#### Methods assessed in non-null simulations

Each of the methods we assessed in simulation was specified by two components: 1) the way in which patches were represented, and 2) the way in which the patch representations were used for case-control analysis.

For (1), we considered the following methods: i) representation of each patch by a 10-dimensional vector obtained by averaging each meta-marker across all pixels in that patch; ii) flattening of each patch, which is a 40×40×10 tensor, into a vector of length 40 · 40 · 10, running PCA on all the patches, and representation of each patch using its value along the top 20 PCs; (iii) representing each patch using the latent representation learned by a simple two-layer convolutional autoencoder; and iv) representing each patch using the latent representation learned by a single instance of VIMA’s variational autoencoder. For method (iv), we set *C*, the number of latent dimensions of VIMA’s autoencoder, to 20, and we reduced the number of epochs of training to 10; we made these changes both to speed up computation in order to allow for larger-scale simulations and to match the smaller size of the simulation dataset and the simpler nature of the simulated case-control signals.

For (2), we considered the following two methods: a) case-control analysis using VIMA’s microniche-based approach as defined above, and b) case-control analysis by clustering the patch representations using leiden clustering with a resolution of 1, computing the relative abundance of each cluster in each sample, performing a T-test for each cluster to detect association between the relative abundance of that cluster and the sample labels, and reporting the minimal P-value obtained across all the clusters with Bonferroni correction for the number of clusters tested.

In **Supplementary Fig. 8** and **Supplementary Fig. 9**, we show results for methods i.a, ii.a, iii.a, iii) a. In **Supplementary Fig. 10**, we show results for methods iv.a versus iv.b. In **Supplementary Fig. 11**, we compare iv.a to alternative versions of the same method but with different values of the parameter *β* defined above, which controls the strength of the variational penalty in VIMA’s variational autoencoder.

#### Figures of merit used to assess non-null simulations

We used three different figures of merit to assess performance in the non-null simulations: 1) power to reject the null hypothesis of no association at level *α* = 0.05, 2) accuracy at identifying the spatial regions of each sample responsible for that sample’s case vs control status, and 3) number of significant microniches detected for methods i.a through iv.a.

For (1), we computed power in the standard way by computing the fraction of simulations in which the null was rejected by each method at each noise level. For (2), we computed accuracy by checking how well the microniche-level scores could be used to identify which patches are truly case-associated, truly control-associated, or truly null. To do this, we computed the average of two different AUROC metrics: the AUROC for using the microniche-level scores to classify layer 2/3 in cases away from the rest of the patches, and the AUROC for using the microniche-level scores to classify layer 2/3 in controls away from the rest of the patches. If all truly case-associated patches are given positive scores, all null patches are given scores of 0, and all truly control-associated patches are given negative scores, then this metric will equal 1. Conversely, if the microniche-level scores are uninformative then this metric will equal 0.5. For methods i.a through iv.a, we used the microniche-level scores produced by the VIMA software; for method iv.b, we assigned to each patch the T-statistic produced from the T-test applied to the cluster containing that patch. For (3), we used the number of microniches passing the FDR 10% threshold as estimated by VIMA; we did not conduct this analysis for method iv.b as this method does not produce a set of FDR-significant patches.

## Data Availability

All data analyzed during this study are available via three previously published articles^19,39,47^.

## Code Availability

An open-source repository containing code for running VIMA is available at https://github.com/yakirr/vima; an open-source repository containing code underlying all figures and tables is available at https://github.com/yakirr/vimapaper; and an open-source repository containing code underlying all simulations is available at https://github.com/yakirr/vimasim.

## Acknowledgements

We thank M. Brenner, G. Eraslan, E. Hodis, D. Kotliar, R. Madhu, J. Pouget, A. Veres, and the Raychaudhuri lab for helpful discussions and feedback. This work is supported in part by the National Institutes of Health (K08AR085165, T32AR007530, UC2AR081023, R01HG013083, R56HG013083, F30AI157385, T32GM144273, T32HG002295) and the Chan-Zuckerberg Initiative. The content is solely the responsibility of the authors and does not necessarily represent the official views of the National Institutes of Health.

