## Supplemental tables and figures for "Powerful and accurate case-control analysis of spatial molecular data"

### Supplementary Information for “Powerful and accurate case-control analysis of spatial molecular data”

#### Contents

**Supplementary Tables** **2**

**Supplementary Figures** **5**

### Supplementary Tables

| Marker | Comments |
|---|---|
| DAPI | nuclear stain |
| CLIC5 | lining fibroblasts |
| CD34 | sublining fibroblasts, endothelial cells |
| HLADR | antigen presentation |
| CD3 | T cells |
| CD90 | sublining fibroblasts, pericytes |
| CD68 | macrophages |

**Supplementary Table 1: The 7 markers profiled in the RA dataset**. For each marker, we include a description of the main associated cell types or states.

| | | | |
|---|---|---|---|
| Vimentin | CD3 | CD137 | CD40 |
| CD19 | CD126 | CD7 | FoxP3 |
| CD31 | CD66 | VCAM-1 | CD127 |
| CD85j | CD69 | CD49f | CD45RA |
| TIGIT | CD134 | TCRyd | Cytokeratin |
| Ki67 | CD8 | CD36 | CD104 |
| CD15 | CD57 | CD16 | Podoplanin |
| HLA-DR | CD56 | CD38 | CD278 |
| CD152 | CD11c | CD1c | CD120b |
| CD21 | HIF-1a | CD123 | CollagenIV |
| CD274 | MMP12 | CD90 | CD117 |
| CD2 | CD54 | CD45 | CD279 |
| CD4 | CD34 | HLA-ABC | CD5 |

**Supplementary Table 2: The 52 markers profiled in the UC dataset**. Hoechst stains and negative control stains are not included in this table.

| | | | | | | |
|---|---|---|---|---|---|---|
| *ADAMTS3* | *COL11A1* | *FRMPD4* | *ITGB8* | *NOS1* | *RBFOX3* | *SOX6* |
| *ADAMTSL1* | *CSMD1* | *GAD2* | *KAZN* | *NOSTRIN* | *RFX3* | *STXBP6* |
| *ANK1* | *CTSS* | *GALNTL6* | *KCNIP4* | *NPAS3* | *RGS12* | *SULF1* |
| *ASIC2* | *CUX2* | *GPC5* | *KCNMB2* | *NPY* | *RGS6* | *SV2C* |
| *ASTN2* | *DACH1* | *GRID2* | *KIAA1217* | *NRG1* | *ROBO1* | *TACR1* |
| *ATRNL1* | *DCC* | *GRIK3* | *KIRREL3* | *NTNG2* | *ROBO2* | *TAFA1* |
| *BTBD11* | *DCLK1* | *GRIN2A* | *L3MBTL4* | *NXPH2* | *RORB* | *TENM2* |
| *CA10* | *DCN* | *GRIN3A* | *LAMA4* | *PALMD* | *RYR3* | *TH* |
| *CACHD1* | *DGKG* | *GRIP1* | *LAMP5* | *PAX6* | *SATB2* | *THEMIS* |
| *CACNA2D3* | *DLC1* | *GRIP2* | *LHX6* | *PDE4B* | *SCUBE1* | *TLL1* |
| *CALB1* | *DLX1* | *GRM7* | *LRP1B* | *PDGFD* | *SEMA3E* | *TMEM132D* |
| *CARTPT* | *EBF1* | *GRM8* | *LRRC4C* | *PDZD2* | *SEMA6D* | *TMEM255A* |
| *CBLN2* | *EGFR* | *HCN1* | *LRRK1* | *PEX5L* | *SLC14A1* | *TNR* |
| *CD22* | *ETNPPL* | *HPSE2* | *LUZP2* | *PLCB1* | *SLC24A2* | *TOX* |
| *CD74* | *EYA4* | *HS3ST2* | *MEIS2* | *PLD5* | *SLC32A1* | *TSHZ2* |
| *CDH6* | *FBXL7* | *HS6ST3* | *MOG* | *PRKG1* | *SLIT3* | *UNC5B* |
| *CHODL* | *FEZF2* | *HTR2A* | *NDNF* | *PRRT4* | *SMYD1* | *VIP* |
| *CLSTN2* | *FGF12* | *HTR2C* | *NFIA* | *PRRX1* | *SNTB1* | *ZMAT4* |
| *CNTN5* | *FGF13* | *ID3* | *NKAIN2* | *PVALB* | *SORCS1* | *ZNF385D* |
| *CNTNAP5* | *FOXP2* | *ITGA8* | *NLGN1* | *RBFOX1* | *SORCS3* | *ZNF804A* |

**Supplementary Table 3: The 140 gene expression markers profiled in the Alzheimer's dataset**. Negative control probes are not included in this table.

| Cell type | Abund(dementia) | Abund(other) | Enrich. | $P$ | $P_{bonf}$ |
|---|---|---|---|---|---|
| Oligodendrocyte | 0.272189 | 0.204545 | 1.33 | 0.000100 | 0.002400 |
| Microglia-PVM | 0.045113 | 0.038217 | 1.18 | 0.051095 | 1.000000 |
| VLMC | 0.034247 | 0.029630 | 1.16 | 0.002900 | 0.069593 |
| L6 IT | 0.150993 | 0.133333 | 1.13 | 0.001400 | 0.033597 |
| OPC | 0.036145 | 0.032000 | 1.13 | 0.034897 | 0.837516 |
| L6 CT | 0.041176 | 0.038462 | 1.07 | 0.225977 | 1.000000 |
| Endothelial | 0.039683 | 0.038462 | 1.03 | 0.444356 | 1.000000 |
| L2/3 IT | 0.006211 | 0.006061 | 1.02 | 0.648035 | 1.000000 |
| L6b | 0.078788 | 0.078534 | 1 | 0.885911 | 1.000000 |
| Astrocyte | 0.112360 | 0.118343 | 0.949 | 0.020998 | 0.503950 |
| Lamp5 Lhx6 | 0.006250 | 0.006623 | 0.944 | 0.121088 | 1.000000 |
| L5 ET | 0.005587 | 0.007299 | 0.765 | 0.000400 | 0.009599 |
| L4 IT | 0.004785 | 0.006410 | 0.746 | 0.000100 | 0.002400 |
| Sst | 0.013333 | 0.018634 | 0.716 | 0.000100 | 0.002400 |
| L5/6 NP | 0.004762 | 0.007246 | 0.657 | 0.000100 | 0.002400 |
| Pvalb | 0.006623 | 0.010870 | 0.609 | 0.001800 | 0.043196 |
| L6 IT Car3 | 0.026519 | 0.064706 | 0.41 | 0.000100 | 0.002400 |
| L5 IT | 0.007463 | 0.025806 | 0.289 | 0.000100 | 0.002400 |
| Pax6 | 0.000000 | 0.000000 | nan | 1.000000 | 1.000000 |
| Vip | 0.000000 | 0.000000 | nan | 1.000000 | 1.000000 |
| Chandelier | 0.000000 | 0.000000 | nan | 1.000000 | 1.000000 |
| Lamp5 | 0.000000 | 0.000000 | nan | 1.000000 | 1.000000 |
| Sncg | 0.000000 | 0.000000 | nan | 1.000000 | 1.000000 |
| Sst Chodl | 0.000000 | 0.000000 | nan | 1.000000 | 1.000000 |

**Supplementary Table 4: Cell types enriched in dementia-associated layer 6 patches compared to non-associated layer 6 patches**. For each cell type, we list its median abundance (as a fraction of all cells) in dementia-associated layer 6 patches, its median abundance (as a fraction of all cells) in the remainder of the layer 6 patches, the enrichment this corresponds to, the P-value generated by randomly switching the associated and non-associated patches within each sample with 10,000 permutations (Methods), and the Bonferroni-corrected P-value.

#### Supplementary Figures

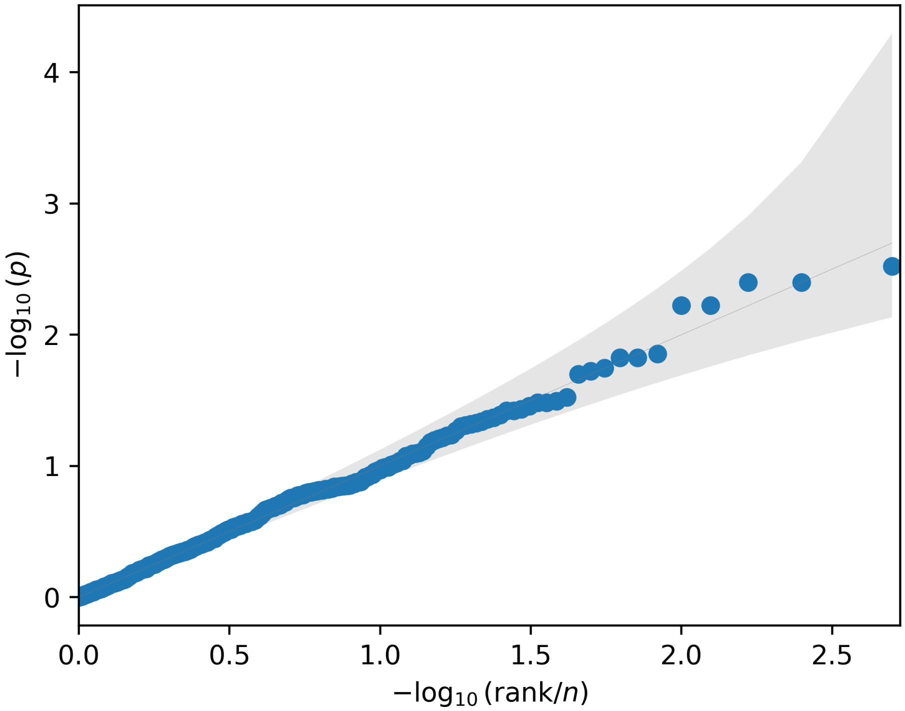


**Supplementary Figure 1: Null calibration of VIMA**. A Q-Q plot comparing the distribution of VIMA global P-values for case-control association obtained across the 500 null trials with a uniform distribution. VIMA's type I error rate across these trials is $27/500 = 0.054 \pm 0.005$.

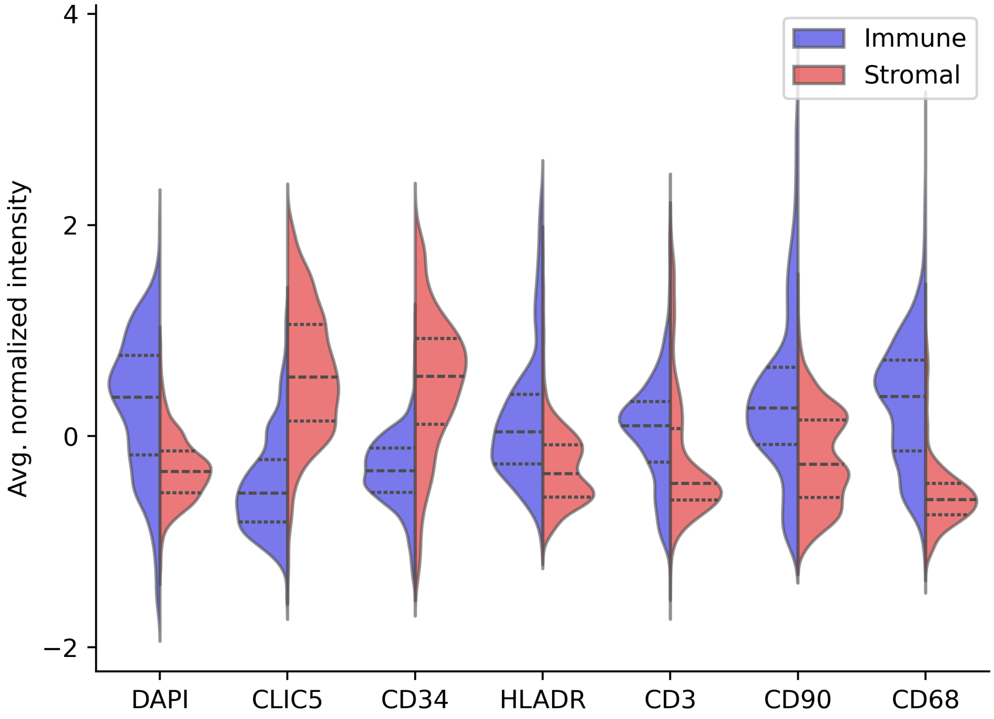


**Supplementary Figure 2: Differentially abundant markers between stromal-associated and immune-associated patches in the RA dataset.** For each of the 7 markers in the dataset, we show the distribution of average marker intensity in patches significantly associated with stromal samples versus those significantly associated with immune samples.

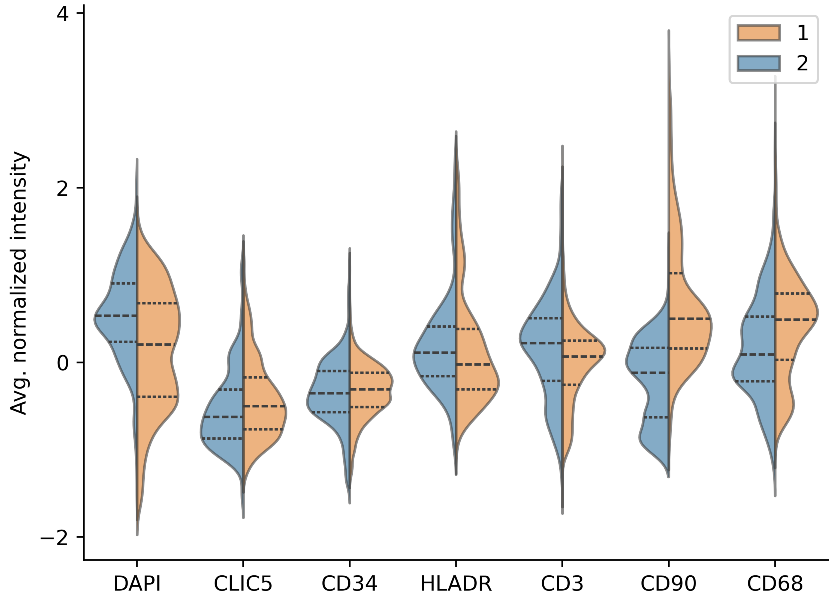


**Supplementary Figure 3: Differentially abundant markers between the two clusters of immune-associated patches in the RA dataset.** For each of the 7 markers in the dataset, we show the distribution of average marker intensity in patches assigned to cluster 1 from Figure 3 versus the average marker intensity in patches assigned to cluster 2 from Figure 3.

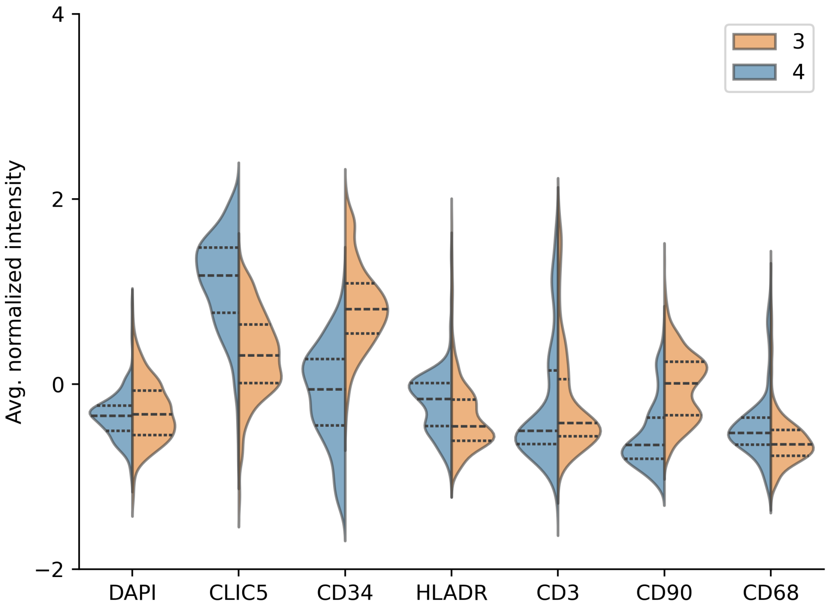


**Supplementary Figure 4: Differentially abundant markers between the two clusters of stromal-associated patches in the RA dataset.** For each of the 7 markers in the dataset, we show the distribution of average marker intensity in patches assigned to cluster 3 from Figure 3 versus the average marker intensity in patches assigned to cluster 4 from Figure 3.

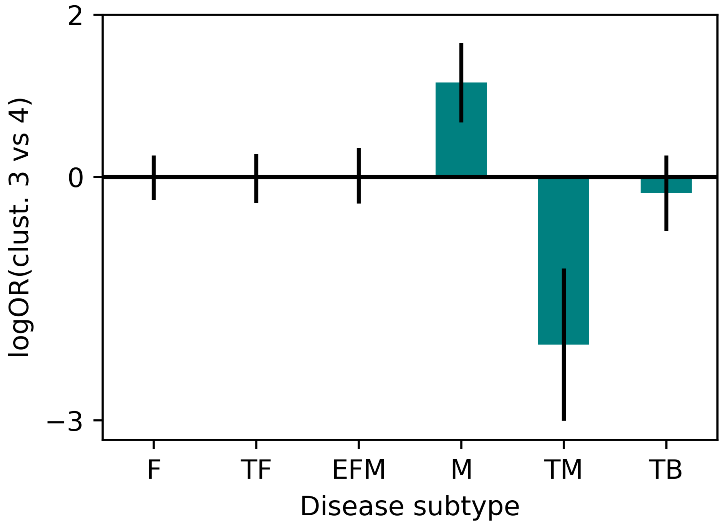


**Supplementary Figure 5: Enrichment of cluster 3 vs cluster 4 among the full set of 6 disease subtypes in the RA dataset.** For each CTAP (disease subtype), we show the log-odds-ratio for a randomly chosen patch from clusters 3 and 4 belonging to cluster 3 (positive log-odds) vs cluster 4 (negative log-odds), with 95% confidence intervals.

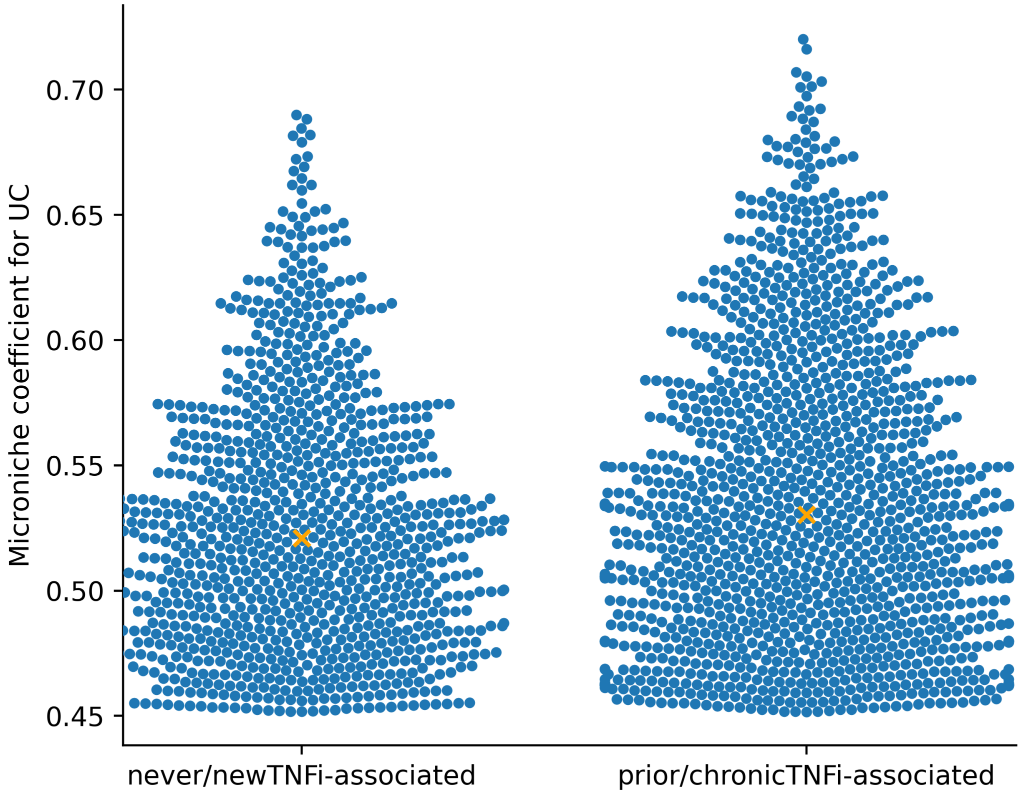


**Supplementary Figure 6: Comparison of UC-vs-control signal to prior/chronicTNFi-vs-never/newTNFi signal.** For all the UC-associated microniches that had a significant correlation to prior/chronicTNFi status, we plot the microniche coefficient for UC status (y-axis) against whether the patch was never/newTNFi-associated (right) or prior/chronicTNFi-associated (left).

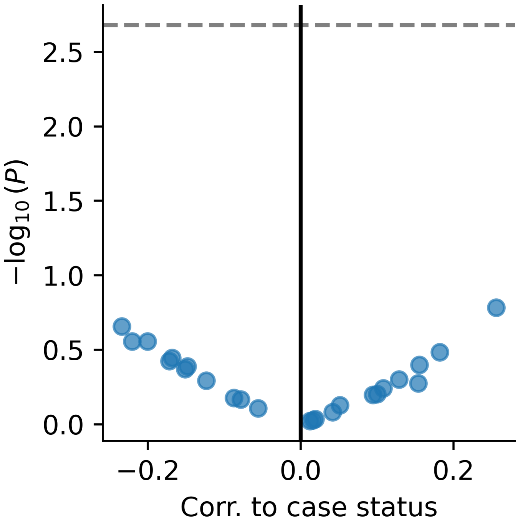


**Supplementary Figure 7: Case-control analysis of Alzheimer's dataset using sample-wide cell-type composition to represent each sample.** We represented each sample using its cell type abundance profile, and then tested each cell type for association and used a permutation-based approach analogous to VIMA's to determine P-values. In the plot, each dot represents a cell type and we plot the correlation of cell-type abundance with case status against the permutation-based P-value for the association. The threshold for significance after Bonferroni correction for the number of cell types tested is shown as a dotted horizontal line.

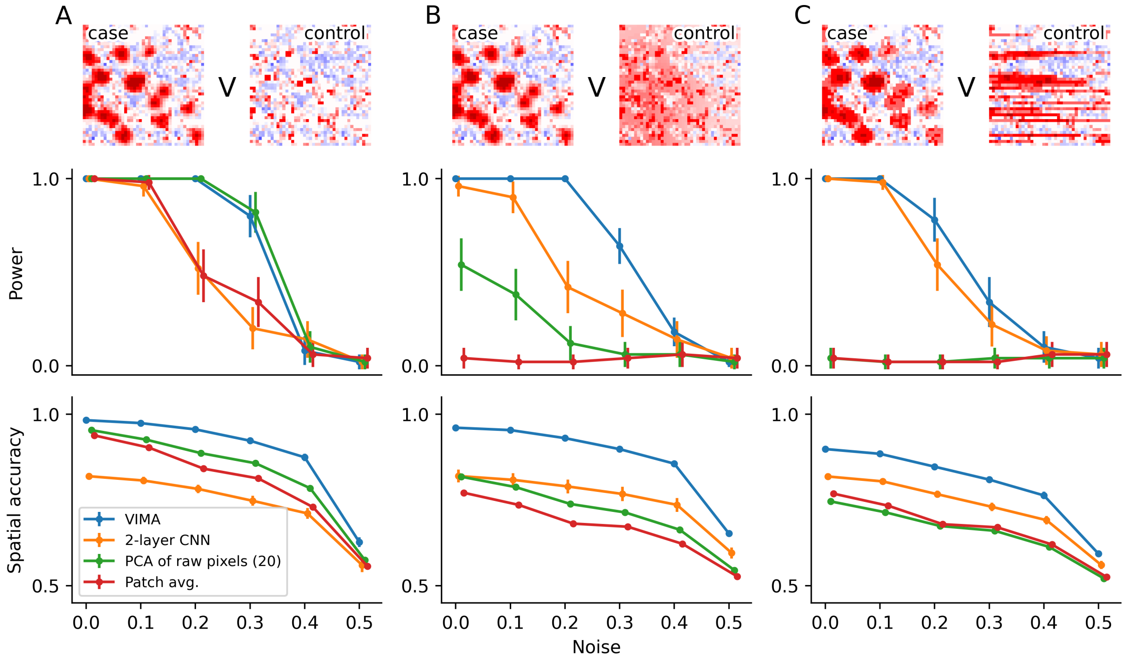


**Supplementary Figure 8: Assessment of increasingly rich patch representations using a real brain spatial transcriptomics dataset augmented with simulated case-control signals.** Each column shows a different signal type spanning A) cellular aggregates added to specific cortical layers in case samples vs no modification to control samples, B) cellular aggregates added to specific cortical layers in case samples vs diffuse infiltrate added to the same cortical layers in control samples, C) cellular aggregates added to specific cortical layers in case samples vs striated tissue structures added to the same cortical layers in control samples. The top row depicts an example tissue patch that has been modified to contain each of the signal types, with the color of each pixel corresponding to intensity of meta-marker 2. The middle row shows the power of each type of patch representation in a single-autoencoder VIMA model at level $\alpha = 0.05$ as a function of increasing amounts of noise. The bottom row shows the spatial accuracy of each method at identifying the regions of each sample that contain the case/control signal, as measured by AUROC (Methods).

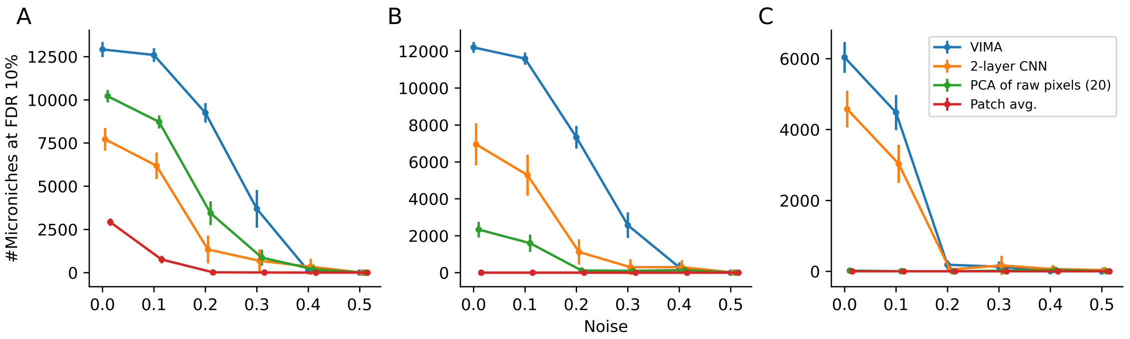


**Supplementary Figure 9: Comparison of number of significant microniches at FDR 10% across signal types and patch representations**. Each plot quantifies the number of microniches achieving significance at FDR 10% as a function of noise for each of the methods shown in Supplementary Figure 8, across the three signal types: A) cellular aggregates added to specific cortical layers in case samples vs no modification to control samples, B) cellular aggregates added to specific cortical layers in case samples vs diffuse infiltrate added to the same cortical layers in control samples, C) cellular aggregates added to specific cortical layers in case samples vs striated tissue structures added to the same cortical layers in control samples.

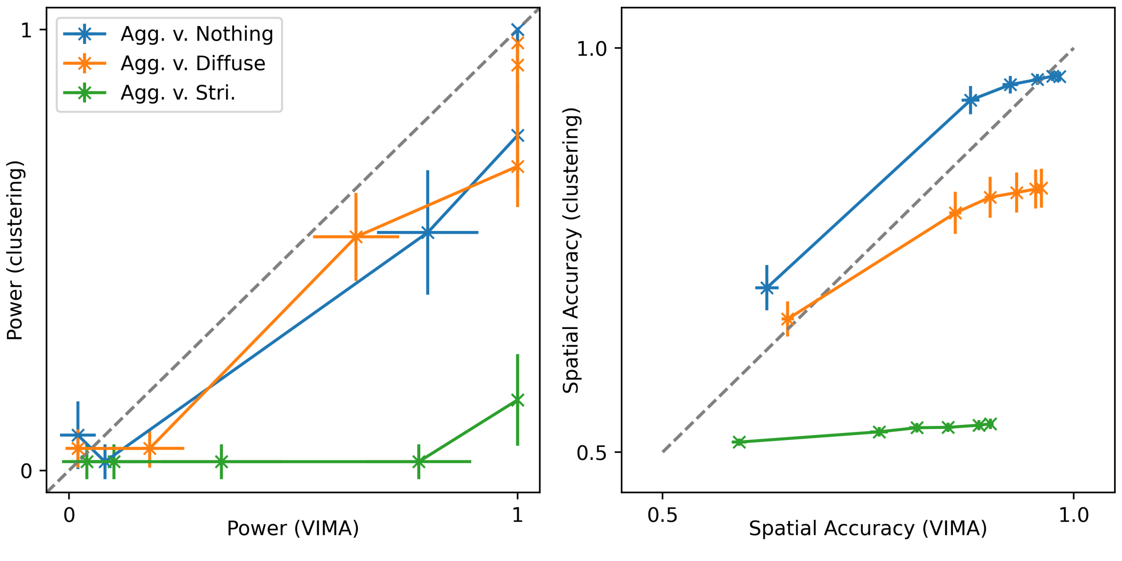


**Supplementary Figure 10: Comparison of microniche- vs cluster-based association testing using VIMA's patch fingerprints**. For each of the three signal types (aggregates versus nothing, aggregates versus diffuse infiltrate, aggregates versus striated structures), and each of 6 increasing noise levels, we show the power (left) and accuracy (right) of the methods in Supplementary Figure 8(x-axis) versus a cluster-based version of each method that performs global and local association testing as in the primary benchmarking study(y-axis).

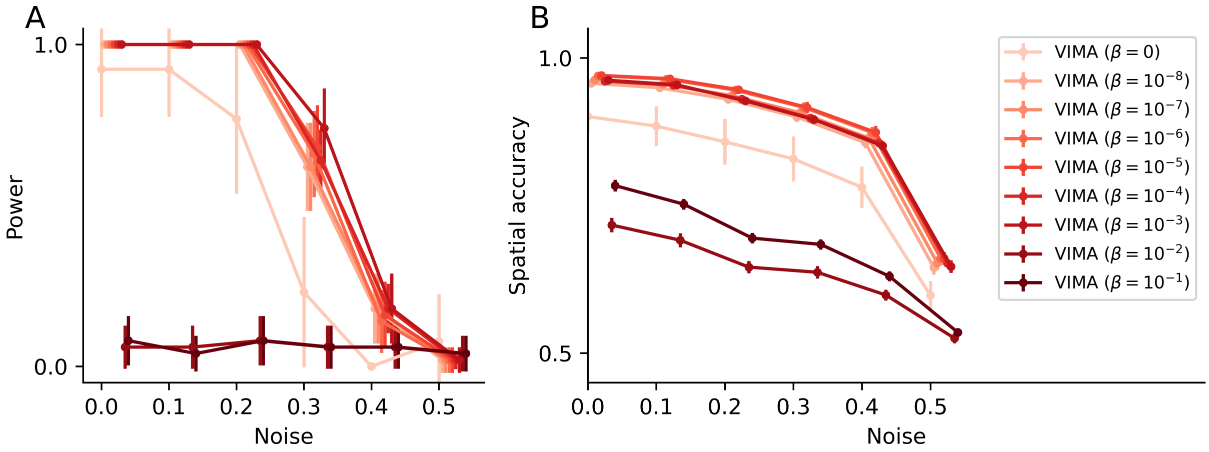


**Supplementary Figure 11: Performance of VIMA with different amounts of variational regularization.** For the cellular aggregates versus diffuse infiltrate signal, we varied the strength of the weight $\beta$ given to the variational penalty in VIMA's VAE across orders of magnitude. Here we show the resulting A) power, and B) spatial accuracy as functions of noise, as in Supplementary Figure 8.

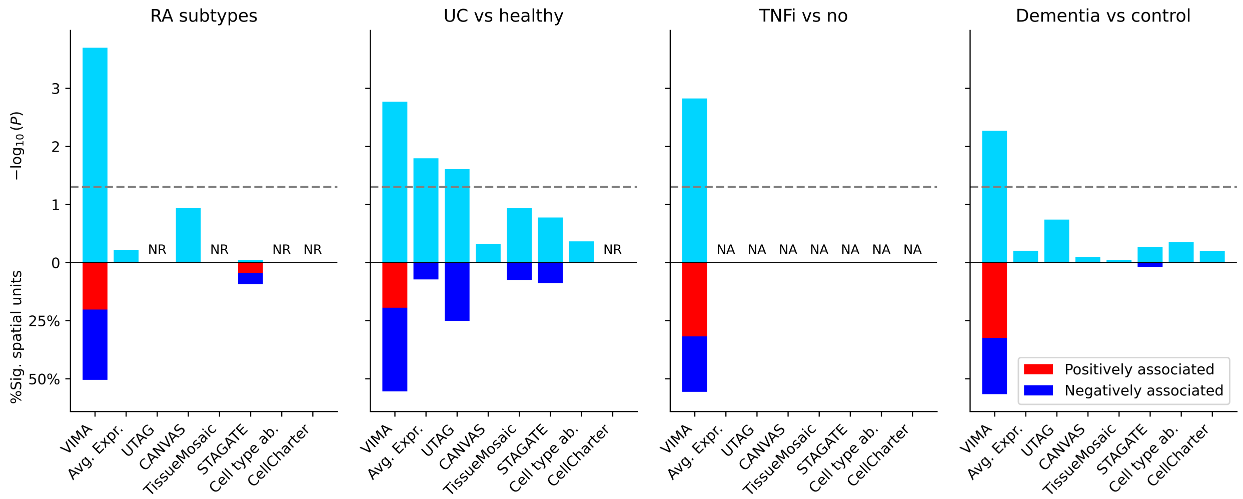


**Supplementary Figure 12: Summary of main benchmarking results.** For each benchmarked method and each of the four phenotypes analyzed in the paper, we plot $-\log_{10}(P)$ for association (upper bar chart) and the fraction of spatial units that passed significance at FDR 10% as estimated using a permutation-based null (bottom bar chart), stratified into positively associated spatial units (red) and negatively associated spatial units (blue). Note that because the FDR estimation framework is not based on a Bonferroni correction, methods with non-significant global P-values can have spatial units that pass FDR significance. Methods that were not runnable on a particular dataset, either because they required cell segmentation (for RA) or did not finish in time (for UC), are labeled as ”NR” (not runnable). For the TNFi analysis, which required UC-associated patches to be run, methods that did not find UC-associated patches are labelled as ”NA” (not applicable).

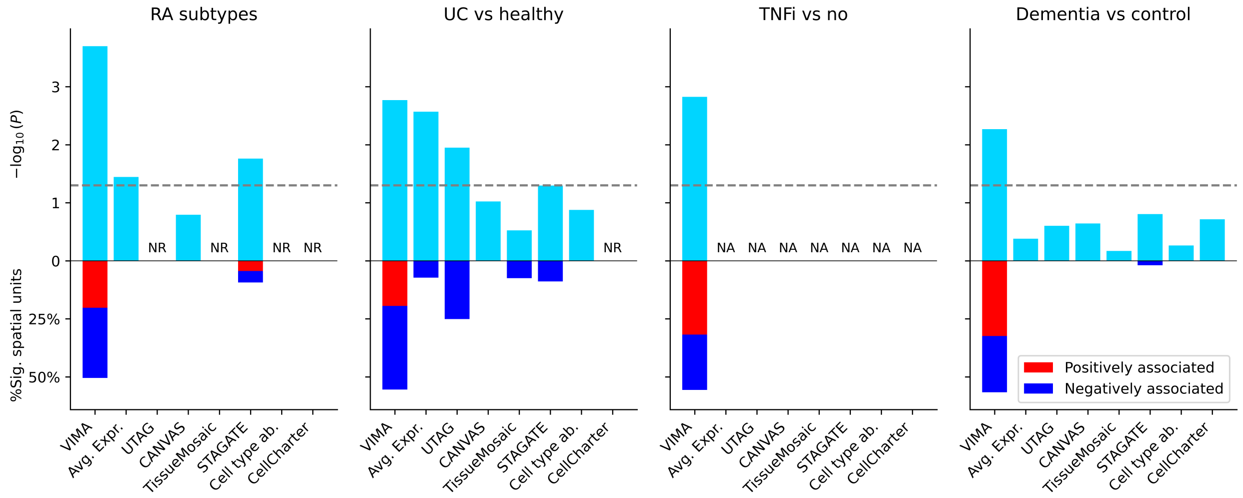


**Supplementary Figure 13: Benchmarking results with VIMA-style global P-values instead of Bonferroni-corrected P-values.** For each non-VIMA method, we conducted the global association test similar to VIMA’s by summing the squared correlations across all clusters and then comparing to a permutation-based null distribution. The significant spatial units are still called using empirical FDR estimation with a cutoff of 10%, as in Supplementary Figure 12. Results are displayed identically to those in Supplementary Figure 12.

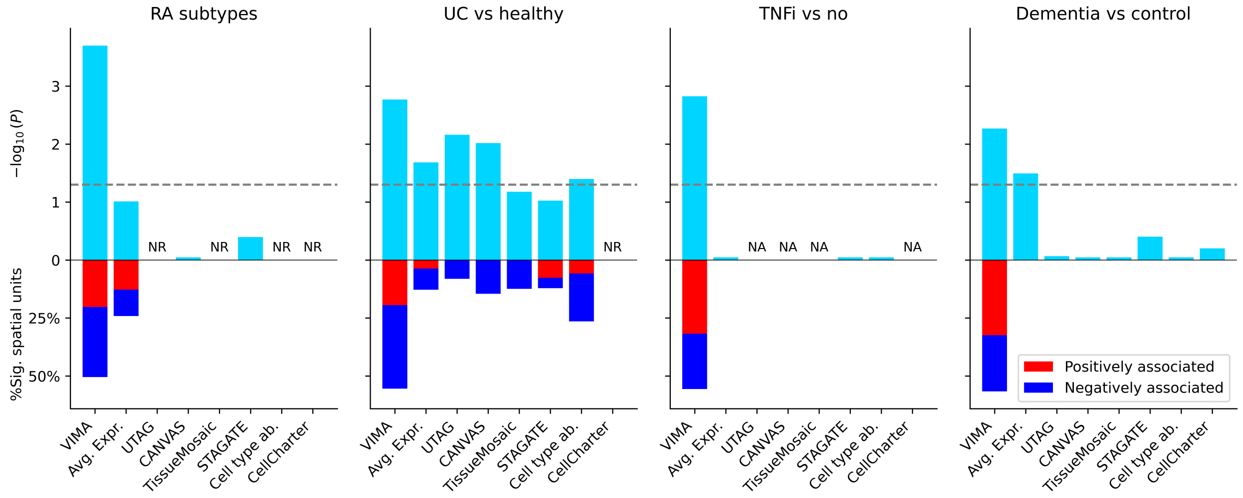


**Supplementary Figure 14: Benchmarking results using post-hoc harmony on tissue embeddings.** For each method, we ran harmony on the resulting tissue embedding prior to clustering and then proceeded as in Supplementary Figure 12. Results are displayed identically to those in Supplementary Figure 12.

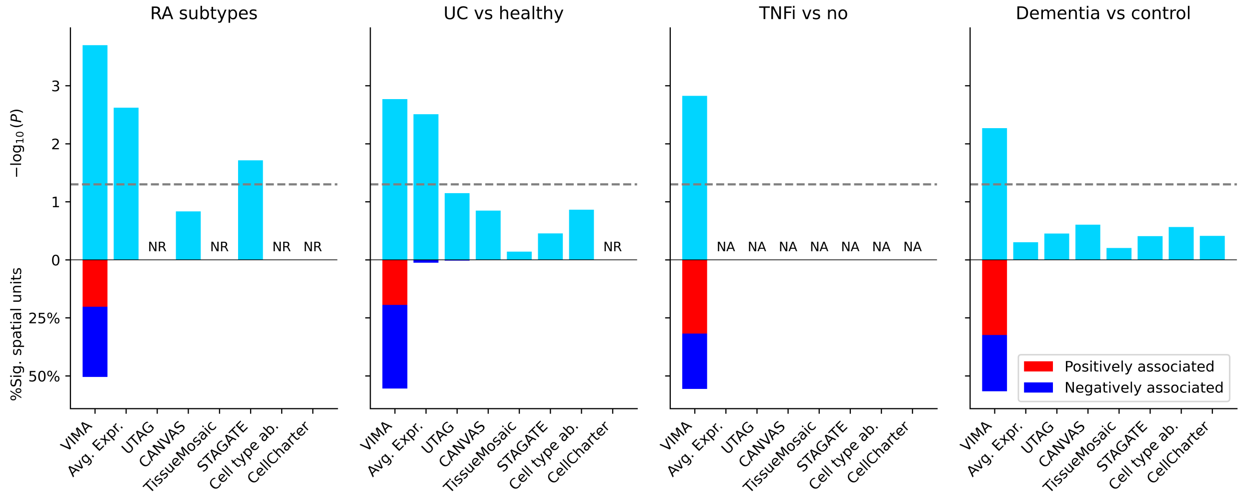


**Supplementary Figure 15: Benchmarking results using microniches instead of clusters.** For each method, we used its tissue embedding to define a set of microniches rather than a set of hard clusters, and fed this representation into the case-control portion of VIMA. Results are displayed identically to those in Supplementary Figure 12.

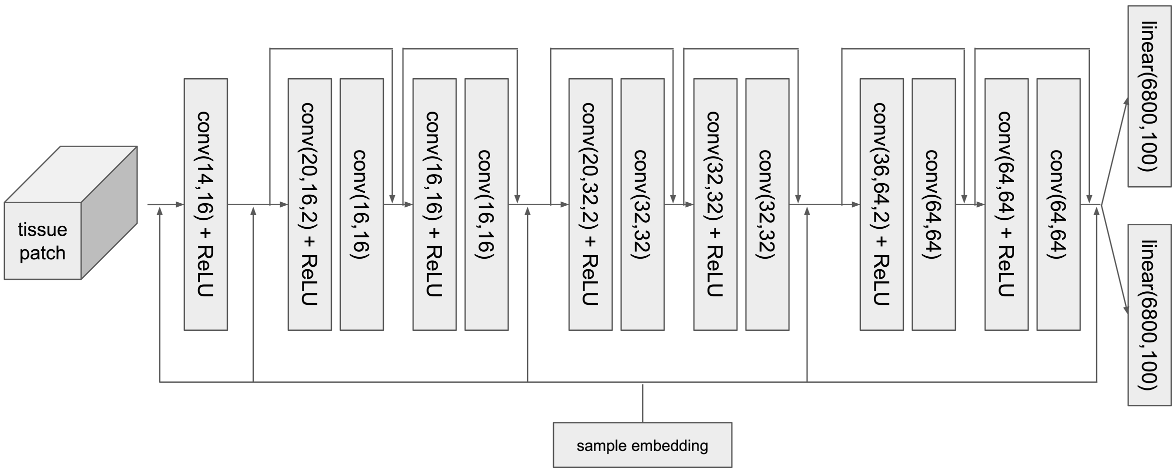


**Supplementary Figure 16: Schematic depiction of the architecture of the VIMA encoder.** The encoder takes as input a tissue patch that by default is of size 40 pixels by 40 pixels by 10 meta-markers. This is passsed through a series of convolutional layers; we denote each convolution by specifying first the number of color channels that it takes as input, second the number of color channels it produces as output, and finally the stride (if it differs from 1). The sample embedding is a 4-dimensional vector that is unique to each sample and learned along with the weights of the model. Each arrow emanating from the sample embedding represents concatenation of these 4 numbers as 4 additional spatially homogeneous color channels. The arrows that begin before and end after some of the convolutional layers represent simple addition, with a convolutional layer to reduce the x- and y- dimensions of the patch if necessary; these “skip” connections, which allow the convolutional layers to learn residual rather than cumulative signal, are the reason for the “Res” in the name “ResNet”.
